# Integrated landscape genomics reveals biogeography and climate-driven local adaptation in high-altitudes

**DOI:** 10.1101/2025.07.02.662798

**Authors:** Nan Wang, Prashant Ghimire, Pritam Chhetri, Nishma Dahal, Cheng Yi, Tong Zhang, Suonan Zhuoga, Zhaxi Jiangyong, Sangeet Lamichhaney

## Abstract

High-elevation environments harbor unique species adapted to altitudinal and environmental extremes. However, due to logistical challenges of sampling such species throughout their distribution across steep and varied terrains, studies often lack comprehensive evidence to understand patterns of population divergence, local adaptation, and how current adaptations may influence future vulnerability. In this study, we utilized Tibetan Partridge *(Perdix hodgsoniae)*, a high-altitude endemic bird found between 2800–5000 meters across the arid western and humid northeastern regions of the Sino-Himalayan landscape. This region’s complex topography-characterized by tall mountains, deep valleys, and contrasting climatic conditions provided opportunities for investigating population divergence, local adaptation, and climate-related vulnerability. We integrated population-scale whole-genome sequencing with ecological, climatic, landscape, and morphological data to examine current patterns of local adaptation and forecast future risks. Our findings show that both biogeographic barriers and climatic variation drive rapid population divergence in *P. hodgsoniae*, reflected in distinct morphological traits and population genetic structure. Western populations, inhabiting dry and fragmented landscapes, exhibit adaptations to temperature extremes with low genetic diversity, reduced habitat suitability, limited gene flow, and weak connectivity—factors that increase their vulnerability to future environmental changes. In contrast, northeastern populations, living in more humid regions, show genetic adaptations linked to precipitation, maintain high genetic diversity and habitat connectivity, and may serve as evolutionary refugia under future climate scenarios. This study underscores the value of integrating genomic, ecological, and landscape data to reveal mechanisms of divergence and adaptation, and to develop robust predictions for conservation planning under rapidly changing environmental conditions.

## Introduction

Mountains are unique ecosystems shaped by complex geological processes. The formation of mountains, deep valleys, rivers, and drainage systems has driven biodiversity diversification. As a result, mountains host approximately 87% of the world’s terrestrial biodiversity within just 25% of the landmass (Colwell et al. 2008; Rahbek et al. 2019). However, these ecosystems are among the most vulnerable to climate change(Knight 2022). Extensive research has been conducted on mountain ecosystems to understand biodiversity patterns (Peters et al. 2019; Rahbek et al. 2019; Hu et al. 2020). Such studies have indicated that geological events, such as plate tectonics and climatic oscillations, including glacial cycles, have left significant imprints on mountain biodiversity (Descombes et al. 2017; Hu et al. 2020). Our current understanding of mountain ecosystems largely comes from broad community-level studies examining changes in species richness, abundance, and diversity along elevational gradients. These studies have demonstrated that many species are shifting toward higher elevations in response to climate change (Freeman et al. 2018; Rödder et al. 2021). However, much less is known about species that already inhabit mountain summits, particularly the endemic species.

High-altitude endemic species are particularly vulnerable to environmental change, as they already face physiological challenges from low oxygen availability and atmospheric pressure (Laguë 2017; Storz 2021). Unlike lowland species that can track their favorable niches by moving upslope in response to changing environment, species confined at the top have limited flexibility to track their favorable niches and hence, face severe constraints (Freeman et al. 2018; Rödder et al. 2021). Their range expansion is restricted not only by existing geographical barriers but also by the stark climatic contrasts within mountain landscapes (Sánchez-Montes et al. 2018). This raises critical questions: How do these species persist in complex biogeographic regions with fragmented habitats and variable environmental conditions? Are their responses to environmental changes uniform across their habitat distribution, or do local biogeography and environmental factors drive distinct adaptive strategies? Additionally, do certain local populations exhibit greater vulnerability due to variations in biogeographic constraints and their genomic architecture?

Addressing these questions requires a comprehensive study of the local populations of such high-altitude endemic species across their distributional range, encompassing diverse landscape and climatic gradients. By integrating landscape and climatic data with population-scale genomic data, we can gain deeper insights into how populations respond to different climatic pressures across their range and identify genetic signatures of local adaptation. Additionally, integrating landscape genomics and eco-evolutionary modeling approaches will allow prediction of “genomic offset”, a measure of the genetic changes required for a local population to adapt to future environmental changes (Rellstab et al. 2021; Chen et al. 2022). Such a comprehensive framework will allow us to accurately identify vulnerable populations and landscapes, enhancing our ability to develop targeted conservation strategies (Bay et al. 2018; Turbek et al. 2023).

The species residing in the Sino-Himalayan region, encompassing the Himalayas and the Qinghai-Tibetan Plateau can offer one such framework to study how species persist in fragmented landscapes and how their genetic adaptations may influence their future survival in a rapidly changing environment. The region’s intricate topography, shaped by the continuous collision of the Indian and Eurasian tectonic plates over the past 50 million years, along with past and ongoing climatic fluctuations, has created substantial biogeographic barriers, including mountain ranges, deep valleys, and expansive plateaus, driving species isolation and diversification (Favre et al. 2015). Recognized as a biodiversity hotspot, this region harbors numerous endemic flora and fauna uniquely adapted to extreme high-altitude environments.

Studying endemic species in the Sino-Himalayan region not only offers insights into historical evolutionary processes but also helps us understand how organisms adapt to ongoing climate change in extreme environments. One such high-altitude endemic species is the Tibetan partridge (*Perdix hodgsoniae*), which inhabits montane shrublands at elevations ranging from 2,800 to 4,600 meters(McGowan and Madge 2010) (Fig. 1a). Although capable of short flights, *P. hodgsoniae* primarily relies on walking, making it largely sedentary with minimal altitudinal migration (Lu and Ciren 2002). This limited mobility, combined with their geographic isolation, results in the formation of multiple local populations across their range.

**Figure 1:**
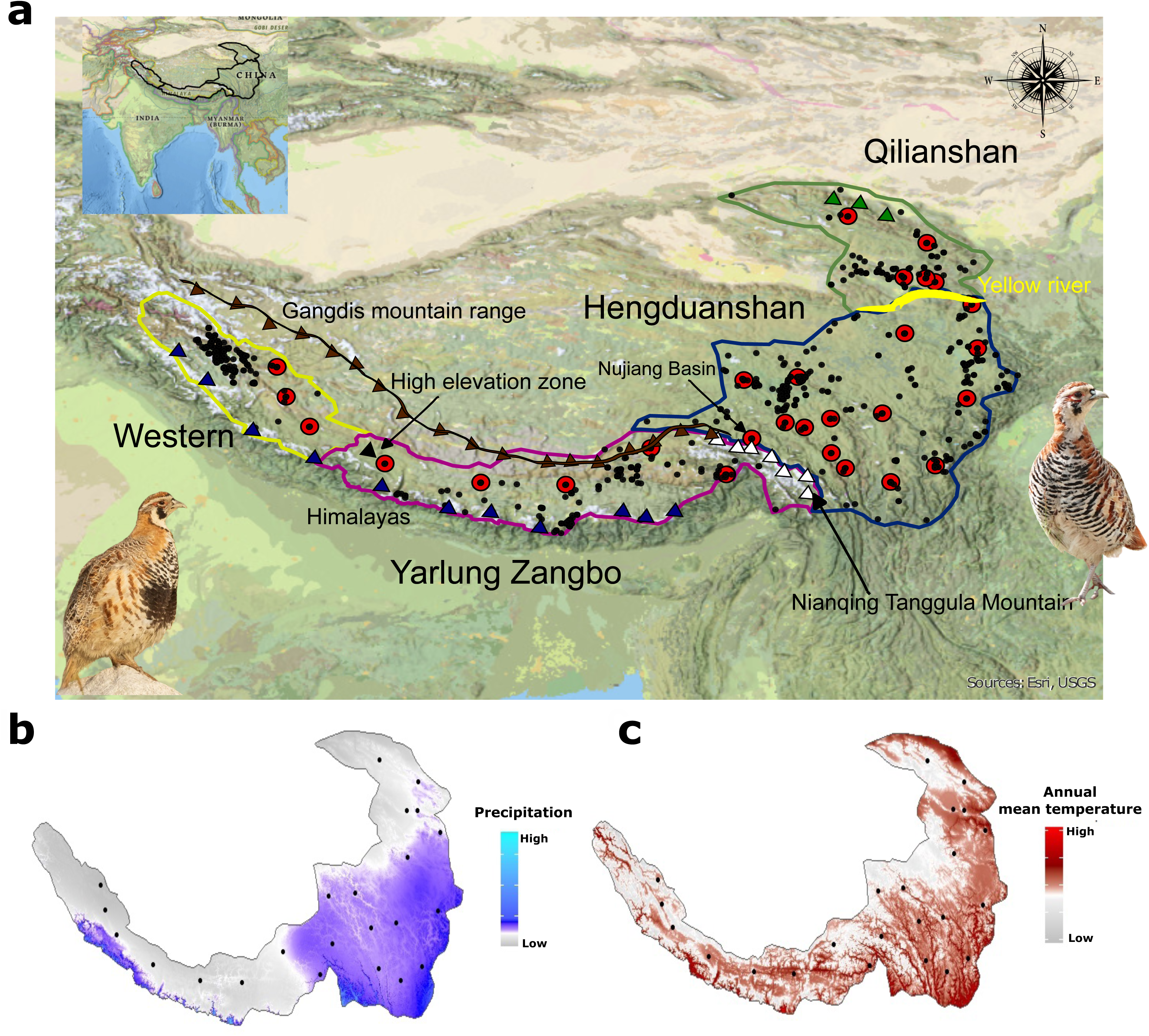
Distribution Range of the Tibetan Partridge and Environmental Variability Across the Sino-Himalayan Landscape. **(a)** The map inset (top left) illustrates the Indian subcontinent for geographical reference. The main panel highlights the distribution range of the Tibetan Partridge, along with key mountains and geographical barriers. Sampling locations are represented by red circumpuncts, while black dots indicate known distribution points based on the eBird database. The high-elevation zone along the upper Yarlung Zangbo River is marked with a black triangle, and prominent mountain ranges include, the Himalayas (blue triangle), Gangdis Mountains (brown triangle with connecting brown line), and Nianqing Tanggula Mountains (white triangle), Qilian Mountains (green triangle). The Yellow River is depicted in yellow. The Tibetan Partridge populations west of the Nianqing Tanggula Mountains are noted for their distinctive black belly patch (bottom left image, © Neeraja V, ML621931282), contrasting with eastern populations that lack this feature (bottom right image, © Nick Addey, ML612172194) **(b)** Annual precipitation (mm) and **(c)** annual mean temperature (x10°C) maps highlight climatic variability across the distribution range.

The western portion of its range, bordered by the Himalayas to the south, the Gangdis Mountains to the north, and the Nianqing Tanggula Mountains in the central east, experiences lower precipitation(Favre et al. 2015) (Fig. 1a, b). In contrast, areas east of the Nianqing Tanggula Mountains, characterized by mid-elevation valleys, receive higher precipitation (Fig. 1a, b). The species’ northern distribution is limited by the Qilian Mountains, with the Yellow River and its tributary, the Huangshui River, forming a natural boundary separating it from the Hengduan Mountains **(Fig. 1a).** Given the substantial variation in precipitation across its distribution range, we hypothesize that precipitation exerts a stronger influence than temperature on local adaptation in Tibetan Partridge populations **(Fig. 1b,c)**. We predict to identify genomic signatures of selection associated with precipitation-related variables, reflecting the orographic effects that drive environmental heterogeneity. Populations in wetter north-eastern regions and drier western regions may exhibit distinct genetic adaptations corresponding to their respective climatic conditions.

In this study, we have developed Tibetan Partridge as a model for understanding how animals with ranges spanning diverse high-altitude habitats may respond to current and future environmental changes. In addition, we have highlighted how integrated analysis of landscape, genomic and climate data can provide a comprehensive understanding of which populations may be more resilient or vulnerable, helping to inform conservation strategies for any species facing global change. We have quantified the patterns of biogeography and climate of the Sino-Himalayan landscape and quantified their impact on divergence and species’ responses to environmental changes. We have further employed ecological and landscape genomics approaches to identify populations that are at greater risk from climate change and map potential gene flow routes that could facilitate their evolutionary rescue.

## Results and Discussion

### Geography and historic isolation shape morphology and population genetic structure in Tibetan Partridges

We measured nine morphological traits from 60 Tibetan partridge individuals sampled across their distribution range in the Qinghai-Tibet Plateau (QTP) region of the Sino-Himalayas (see methods for details). Our analysis revealed notable morphological variation among local populations of Tibetan partridge. Beak, paw, and tarsometatarsus length were significantly different across landscapes (p < 0.05), with larger size in the north-eastern populations (from Hengduanshan and Qilian landscapes), while wing and tail length showed little variation across the landscape but tended to increase from the western (from Western and Yarlung Zangbo landscapes) to the northeastern regions **(Fig. 2a, Supplementary Fig. 1a-h).** To determine whether the morphological traits exhibited distinct patterns across each landscape, we performed a principal component analysis (PCA) on these nine morphological traits that revealed a clear separation between the western and north-eastern populations, indicating morphological differences among local populations of Tibetan Partridges residing in these different habitats **(Fig. 2b)**. Our morphological analysis indicates that Tibetan partridge populations in dry landscapes tend to have smaller body sizes compared to those in wetter environments. This difference likely reflects the landscape features such as mid-elevational region on wet Hengduanshan, and birds’ reliance on shrubland for food, which is relatively sparse in the drier western regions **(Fig. 1c, Supplementary Fig. 2a,b)**, and smaller body size offering an adaptive advantage in these resource-limited conditions. In contrast, in wetter regions with abundant vegetation and more readily available food resources, a larger body size may enhance the birds’ ability to efficiently access and store these resources.

**Figure 2:**
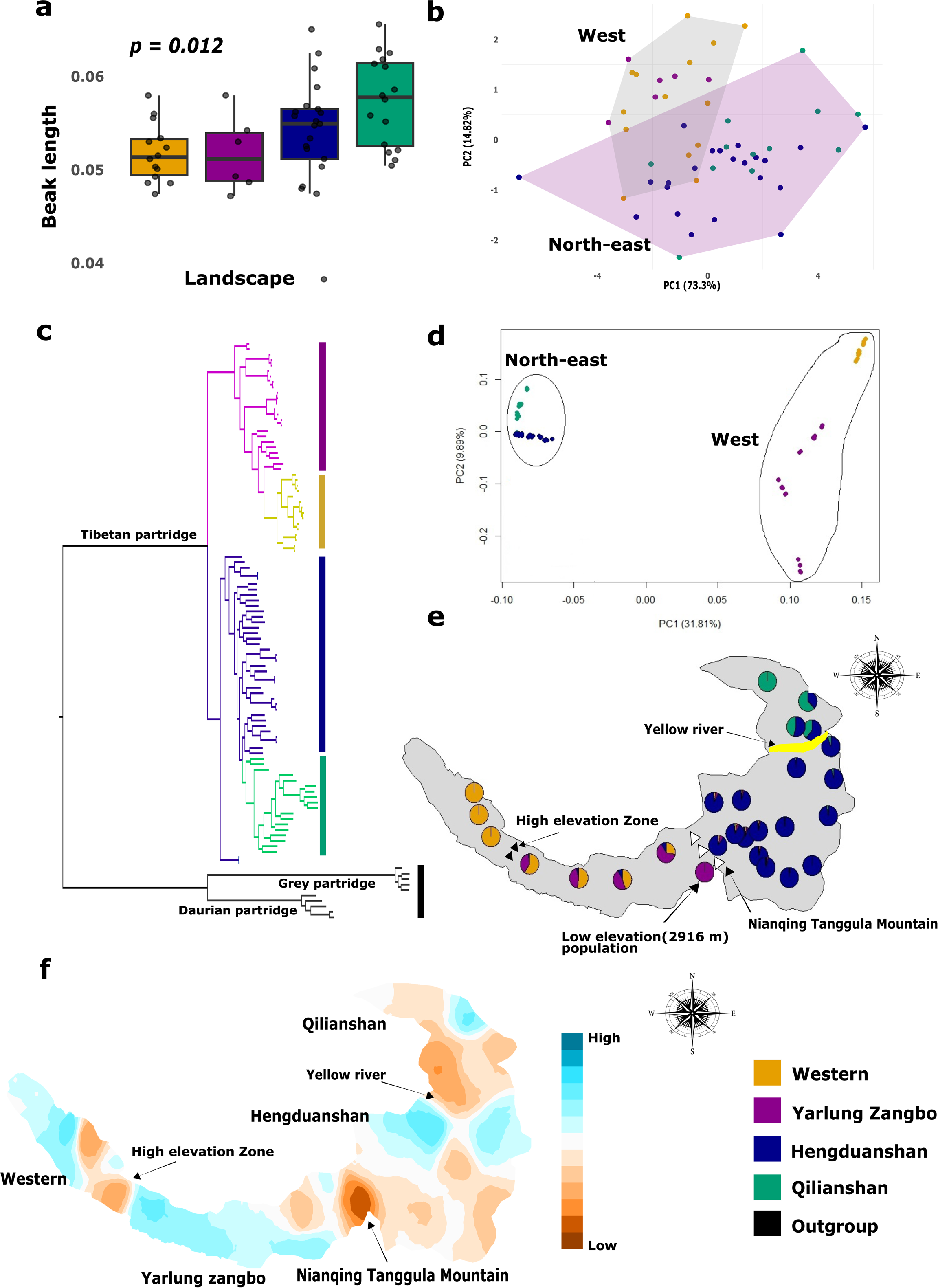
Morphological and Genomic Evidence Reveal Geographic and Historical Isolation as Key Drivers of Genetic Structure in Tibetan Partridges. **(a)** Beak length variation: adjusted beak length (mm) by body weight across landscapes shows an increasing trend from the Western landscape to the Qilianshan region **(b)** Morphological PCA: Principal Component Analysis (PCA) of nine morphological traits indicates two distinct clusters, reflecting variation across Tibetan Partridge populations **(c)** Phylogenetic analysis: A maximum likelihood phylogenetic tree constructed from 29,595,022 SNPs identifies the genetic structure of Tibetan Partridge populations, strongly aligned with their respective landscapes. All major nodes have local support > 0.9 based on the Shimodaira–Hasegawa test(Price et al. 2010) **(d)** Genetic PCA: PCA based on genomic data reveals clustering patterns within populations, consistent with the phylogenetic tree results, further validating the population structure **(e)** Admixture proportions map: A distribution map displays the admixture proportions for individuals assigned to each genetic cluster (K=4). Key geographical features are annotated, including the high-elevation zone along the Yarlung Zangbo River (black triangles), the Nianqing Tanggula Mountains (white triangles), and the Yellow River (yellow). This map was generated using the Mapmixture v.0.1.061(Jenkins 2024) **(f)** EEMS Analysis: The Estimated Effective Migration Surface (EEMS) illustrates migration rates with color contours: orange indicates low migration rates, while blue signifies high migration rates. Color codes for each landscape are consistent across all panels for clarity and comparison.

We further conducted whole-genome sequencing of 96 individuals of Tibetan Partridges at ∼20-25X average sequence coverage for each sample **(Supplementary Fig. 3)** from 28 different locations across these four landscapes **(Fig. 1a)** and 10 individuals from two closely related species residing at lower elevations; Daurian Partridge *(P. dauurica)* and Grey Partridge *(P. perdix)*. We used this genomic data to generate ∼5.1 million high-quality SNPs that were polymorphic in at least one population. A maximum likelihood phylogenetic tree based on these SNPs identified that populations from each of the four landscapes in western (Western and Yarlung Zangbo) and north-eastern (Hengduanshan and Qilianshan) region formed genetically distinct clades (**Fig. 2c)**. We estimated that the divergence of Tibetan partridge populations occurred between 0.33 (0.35) and 0.05 (0.004) million years ago (mya) **(Supplementary Fig. 4a)**. The initial divergence took place between the western and north-eastern populations around 0.33-0.35 mya, followed by the split of the Nanjiang Basin population from the Qilianshan and Hengduanshan populations in the north-east approximately 0.09-0.11 mya. A subsequent divergence occurred within a similar timeframe (∼0.05-0.07 mya), separating the Qilianshan and Hengduanshan populations. Principal component analysis (PCA) using 291,338 Linkage-Disequilibrium (LD)-pruned SNPs revealed clustering patterns consistent with the phylogenetic tree, clearly separating populations from the western and northeastern landscapes **(Fig. 2d).**

These results indicated that the population divergence of Tibetan Partridge across their distribution range in the Sino-Himalayan landscape is strongly influenced by the region’s mountain ranges and river basins, highlighting the significant role of geography in shaping their genetic divergence.

Admixture analysis revealed distinct population structures at (a) both the western and north-eastern edges of the distribution, (b) in the low-elevation population (2,900 m) within the Yarlung Zangbo landscape and (c) populations from the Hengduanshan landscape **(Fig. 2e)**. In contrast, other populations from the Yarlung Zangbo and Qilianshan landscapes displayed relatively high levels of admixture, suggesting greater gene flow among the populations within these regions. We also employed Estimated Effective Migration Surfaces (EEMS) analysis (Petkova et al. 2016) to map corridors and barriers to gene flow across these landscapes. EEMs analysis identified the Nianqing Tanggula mountains, separating the habitats of western and north-eastern populations, as a potential barrier to gene flow **(Fig. 2f).** It also indicated that high elevation areas at the upper stretches of the Yarlung Zangbo region and north of the Yellow River could act as a barrier to gene flow. Consistent with the results from admixture analysis **(Fig. 2e)**, EEMS highlighted possible gene flow corridors within the Yarlung Zangbo landscape. Additionally, the Hengduanshan region displayed both areas (a) supporting lower gene flow (on the northeast and eastern edges) and (b) regions supporting higher gene flow (to the west and east of the Yellow River) **(Fig. 2f)**. To further characterize the patterns of genetic variation across the distribution range of Tibetan partridges, we used Multiple Regression on Distance Matrices (MRM) to assess the relative contributions of geographic distance (Isolation by Distance, IBD) and environmental differences (Isolation by Environment, IBE) to genetic differentiation (FST). Our analysis revealed that environmental istance (β = 0.0771, p = 0.001) has a stronger influence on genetic variation compared to geographic distance (β = 0.0000433, p = 0.125), which was not statistically significant. The full model explained 38.1% of the genetic variation (R² = 0.381, p = 0.001), indicating that environmental factors (IBE only, R² = 0.362, p = 0.001) play a more prominent role than geographic distance (IBD only, R² = 0.147, p = 0.002) in shaping the genetic structure of Tibetan partridges. **(Supplementary Fig. 4b,c)**.

Phylogenetic dating, admixture, and EEMS analyses indicate that the biogeographic history of the Tibetan Partridge has been shaped by three major geographic barriers, resulting in the formation of genetically distinct local populations. The primary barrier is the Nianqing Tanggula Mountains, which separated western and northeastern populations approximately 0.35 million years ago. Subsequently, the Yellow River isolated the Hengduanshan population around 0.06 million years ago, while high-elevation zones along the upper Yarlung Zangbo River further separated the western and Yarlung Zangbo populations approximately 0.06 million years ago. It is unlikely that these divergence events were driven by the uplift of the Qinghai-Tibetan Plateau (QTP), as the region had already reached its current altitude by the late Pliocene (2.58 million years ago) (Wang et al. 2014; Deng and Ding 2015). Instead, similar to the populations of Snow Partridge, which is previously known to diverged during the same period (∼0.4 million years ago) in QTP (Yao et al. 2022), early divergence in Tibetan Partridges may have also been influenced by interglacial isolation. Notably, the latter two divergence events occurred almost simultaneously during Marine Isotope Stage (MIS) 3 (0.06–0.03 million years ago), a period of extensive glaciation in the QTP that caused significant climatic fluctuations(Owen et al. 2003; Cui et al. 2018). While previous studies have highlighted the role of the Nianqing Tanggula Mountains in shaping the west and north-east divergence of diverse species in the Sino-Himalayan region(Liu et al. 2018; Du et al. 2020; Rana et al. 2021), the impact of high-elevation zones in the upper stretches of the Yarlung Zangbo and the Yellow River had remained largely unexplored. Earlier studies from thisin this region were constrained by limited sampling across species distribution range (especially from Tibet)(Du et al. 2020; Chen et al. 2022; Chen et al. 2023; Jiao et al. 2024). This study presents a comprehensive sampling of Tibetan partridges across these regions, offering new insights into their biogeographic history and the influence of diverse landscapes on population divergence.

### Tibetan partridge populations demonstrate genetic adaptation to local climatic conditions

Climatic variation in the Sino-Himalayan landscape has led to the population divergence of many species in the region(Yao et al. 2022). There is a considerable variation in temperature and precipitation across different landscapes **(Fig. 1b,c)**, driven by the region’s complex topography, characterized by tall mountain ranges and diverse altitudinal gradients. We expect these differences in bioclimatic variables are crucial for shaping the ecological dynamics, habitat suitability, and adaptive strategies of local Tibetan Partridge populations residing across these varied landscapes. Our comprehensive sampling of the local populations of Tibetan partridges across diverse landscapes within the Sino-Himalayan region provided a unique opportunity to study their adaptation to varying local climatic conditions. To identify key bioclimatic variables for Genotype-Environment Association (GEA) analysis, we ranked the importance of all 19 bioclimatic variables available in the WorldClim database from this region (Fick and Hijmans 2017) using the gradientForest analysis (Ellis et al. 2012). Additionally, we calculated pairwise correlations between these variables to ensure the selection of independent bioclimatic variables. From this analysis, we selected five key variables (BIO2, BIO8, BIO9, BIO12, and BIO15) that ranked high in importance, showed low correlation (|r| < 0.6), and represented both temperature and precipitation-related variables **(Supplementary Fig. 5, Supplementary Table 1).**

To identify genomic variants associated with the selected bioclimatic variables, we employed two distinct approaches to minimize false positives. First, we used the Latent Factor Mixed Model (LFMM) (Frichot et al. 2013) to examine univariate relationships between SNPs and each bioclimatic variable. The LFMM analysis identified 442,926 candidate SNPs associated with these five bioclimatic variables after adjusting for multiple testing using FDR correction (p < 0.05) **(Fig. 3a, c & Supplementary Fig. 6a-c).** Candidate SNPs associated with specific bioclimatic variables are expected to exhibit significant allele frequency differences among individuals residing across these climatic gradients. To evaluate this, we calculated pairwise genetic divergence (FST) between 10 individuals from habitats with the highest and lowest values for these bioclimatic variables. The analysis confirmed that the candidate SNPs showed significantly greater genetic differences compared to an equal number of randomly selected SNPs **(Supplementary Fig. 7a-e)**, aligning with our prediction. Most (94.96%) of the candidate SNPs identified by LFMM univariate analysis were associated with precipitation-related variables (BIO12 and BIO15), whereas only relatively smaller proportions (5.04%)) were associated with temperature-related variables (BIO2, BIO8, and BIO9) **(Supplementary Fig. 8)**. These results support our hypothesis that precipitation plays a stronger role than temperature in driving local adaptation in Tibetan Partridge populations.

**Figure 3:**
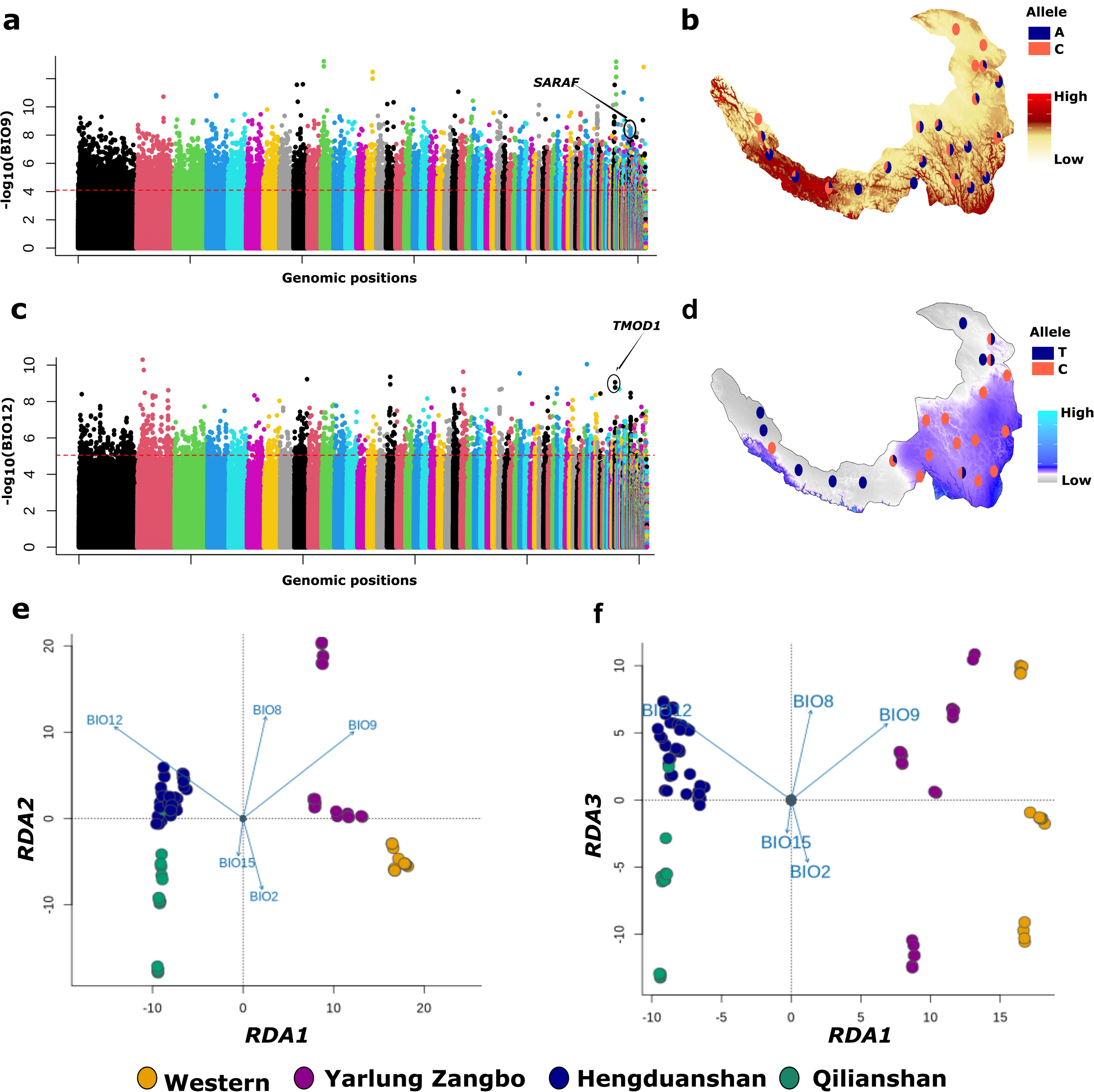
Genetic Adaptations of Tibetan Partridge Populations to Local Climatic Conditions. **(a)** Manhattan plot showing -log10(p) values from the Gene-Environment Association (GEA) analysis for 5,068,604 SNPs with BIO9 (Mean Temperature of the Driest Quarter) as the environmental variable. Candidate SNPs surpassing the red dashed line meet the 1% False Discovery Rate (FDR) correction threshold. The *SARAF* gene (*Store-operated calcium entry-associated regulatory factor*) overlapping one of the top candidate SNP, is highlighted within a circle **(b)** The allelic distribution of a candidate SNP (*Scaffold_631:373769*) within the *SARAF* gene demonstrates a higher frequency of the C allele in populations inhabiting regions with lower mean temperatures during the driest quarter (BIO9) **(c)** Manhattan plot showing -log10(p) values for SNPs associated with BIO12 (Annual Precipitation). Candidate SNPs surpassing the red dashed line meet the 1% False Discovery Rate (FDR) correction threshold. The *TMOD1* gene (*Tropomodulin-1*), overlapping one of the top candidate SNP, is highlighted within a circle **(d)** The allelic distribution of a candidate SNP (*Scaffold_720:849107*) within the *TMOD1* gene reveals contrasting patterns: the T allele is nearly fixed in drier regions (white), while the C allele dominates in wetter regions (blue) (e) Redundancy analysis showing results of the first two axes **(e)** and first and third axis **(f)**; Each dot represents an individual bird, color-coded by its population landscape (consistent with Fig. 1). Blue vectors represent the direction and magnitude of correlation with environmental predictors represented by five composite bioclimatic variables principal component. Populations from Hengduanshan and Qilianshan landscapes are associated with precipitation-related variables (Annual Precipitation: BIO12 and Precipitation Seasonality: BIO15), while populations from Western and Yarlung Zangbo landscapes are associated with temperature-related variables (Mean Temperature of Driest Quarter: BIO9, Mean Temperature of Wettest Quarter: BIO8, and Mean Diurnal Range: BIO2).

The second approach we used for GEA analysis was multivariate redundancy analysis (RDA) (Capblancq and Forester 2021) to identify genomic regions associated with previously selected five bioclimatic variables (BIO2, BIO8, BIO9, BIO12, and BIO15). The first three RDA components explained, respectively, 66.76%, 17.47% and 6.18% of the variation. The results showed that individuals residing in the same landscapes clustered together, indicating exposure to similar climatic conditions within each landscape **(Fig. 3e,f)**. Both RDA plots, RDA1 vs. RDA2 **(Fig. 3e)** and RDA1 vs. RDA3 **(Fig. 3f)** revealed that genetic variation in individuals from Hengduanshan and Qilianshan landscapes are positively related with high Annual Precipitation (BIO12) and Precipitation Seasonality (BIO15). In contrast, genetic variation in individuals from the Western and Yarlung Zangbo landscapes are positively related with Mean Diurnal Range (BIO2), Mean Temperature of Wettest Quarter (BIO8), and Mean Temperature of Driest Quarter (BIO9), demonstrating varying climatic associations between the western and north-eastern populations of Tibetan partridges. These results, indicate distinct climatic associations among Tibetan Partridge populations across their distribution range, aligning with our hypothesis. While precipitation-related variables predominantly drive genetic variation in the northeastern populations, temperature plays a more significant role in shaping genetic variation in the Yarlung Zangbo and western landscapes. In dry regions, species must adapt to extreme heat and limited water availability, making temperature regulation crucial for survival (Fuller et al. 2021). Conversely, in wetter north-eastern regions, adaptation to high moisture levels and excessive precipitation is the key. These findings underscore how diverse environmental factors across heterogeneous landscapes drive divergent evolutionary responses, shaping local adaptations in Tibetan Partridges.

We further performed outlier detection using redundancy analysis and identified 14,999 candidate SNPs (0 SNPs RDA1, 164 SNPs RDA2, 14835 SNPs at 3.5 SD) associated with these five bioclimatic variables (BIO2, BIO8, BIO9, BIO12, and BIO15). RDA1 is most strongly associated with BIO12 (Annual Precipitation; loading = -0.67) and BIO9 (Mean Temperature of Driest Quarter; loading = 0.58), while RDA2 is primarily driven by BIO8 (Mean Temperature of Wettest Quarter; loading = 0.78) and BIO12 (loading = 0.70). RDA3 shows moderate associations with BIO2 (Mean Diurnal Range; loading = 0.25) and BIO8 (loading = -0.49), though with weaker overall correlations. Of these, 4,377 candidate SNPs were detected by both the univariate (LFMM) and multivariate (RDA) analyses **(Supplementary Fig. 8)**. We defined these overlapping 4,377 SNPs as "climate-associated core candidate SNPs” and were selected for downstream functional annotation and enrichment analysis. One of the top candidate SNPs associated with BIO9 (Mean Temperature of the Driest Quarter) (Scaffold_631:373,769, A/C, p < 0.05) overlaps with the *Store-operated calcium entry-associated regulatory factor (SARAF)* gene. This SNP shows a higher frequency of the A allele in the regions with high mean temperatures **(Fig. 3b)**. *SARAF* activates the mechanistic target of the Rapamycin (mTOR1) pathway, which is linked to heat stress and impacts signal transduction in avian skeletal muscle (Sanlialp et al. 2020). Similarly, another top candidate SNP associated with BIO12 (Annual Precipitation) (Scaffold_720:849,107, T/C, p < 0.05) overlaps with the *Tropomodulin-1 (TMOD1)* gene. This SNP demonstrates near fixation of two alleles in the regions characterized by either dry or wet conditions, highlighting distinct allele frequencies in local populations residing in habitats with varying precipitation levels **(Fig. 3d)**. *TMOD1* which is critical for cellular stability and cytoskeletal organization (Yamashiro et al. 2012), may play a role in cellular responses to osmotic stresses, such as varying precipitation levels.

The majority of these climate-associated core candidate SNPs were non-coding (n = 4,313, 98.5%), and only 64 (1.5%) were coding SNPs **(Supplementary Table 2).** We further extracted 295 genes within the 100 kb region upstream and downstream from these SNPs (to extract their potential regulatory regions) and annotated their Gene Ontology and associated metabolic pathways (**Supplementary Resource 1).** Enrichment analysis of these genes identified significant enrichment of three Gene Ontology terms; Adult behavior (*GO:0030534*, p-value=3.9 x 10^-6^), neurotransmitter receptor internalization (*GO:0099590*, p-value=1.7 x 10^-05^) and diacylglycerol kinase activity (*GO:0004143*, p-value=1.9 x 10^-05^). In addition, we identified 27 missense mutations among these 4,377 “climate-associated core candidate SNPs” spanning 22 genes (**Supplementary Resource 2)**. Although no significant pathway enrichment was detected, these genes include notable examples linked to adaptation to precipitation and temperature. For instance, the *Ras and Rab Interactor 2 (RIN2)* gene is associated with plastic responses to dry environments (Kordonowy and MacManes 2017), while the *Epidermal Growth Factor Receptor (EGFR)* and *Ecto-NOX Disulfide-Thiol Exchanger 2 (ENOX2)* genes are known for their roles in responding to heat stress (Kaushik et al. 2016; Wang et al. 2022). These results indicate that multiple genes and regulatory pathways associated with response to environmental stimuli (Verburg-van Kemenade et al. 2017; Stonehouse et al. 2024) contribute to the climate-driven local adaptation in Tibetan Partridges.

### Genomic offsets and niche modeling reveal vulnerable Tibetan partridge populations across the Sino-Himalayan landscape

The heterogeneous landscape of the Sino-Himalayan region includes diverse environmental and topographical conditions **(Fig. 1b,c, Supplementary Fig. 2a,b)**, which expose local populations of Tibetan Partridges to varying levels of vulnerability to short and long-term environmental changes. We calculated genomic offsets using 4,377 climate-associated core candidate SNPs among the local populations of Tibetan Partridge across their distribution range. Genomic offset measures the genetic distance between current and future genomic compositions, with the latter predicted based on current genotype-environment relationships (Rellstab et al. 2021). A higher genomic offset indicates a greater challenge for a local population to adapt to future climatic scenarios and become more vulnerable to future environmental changes. We employed Gradient Forest (GF) analysis to estimate genomic offset by modeling allele frequency changes along the environmental gradients (Ellis et al. 2012). GF analysis predicted an increase in genomic offset over time (2070 to 2090) and under more extreme climatic conditions **(Supplementary Fig. 9a-c)** among the local populations. At the landscape scale, higher genomic offsets were predicted at the edges of the Tibetan partridge’s distribution range **(Fig. 4a-b, Supplementary Fig. 9a-c).** The western landscape showed the highest genomic offset, followed by Qilianshan, and Yarlung Zangbo, and the lowest genomic offset was predicted for the Hengduanshan landscape.

**Figure 4:**
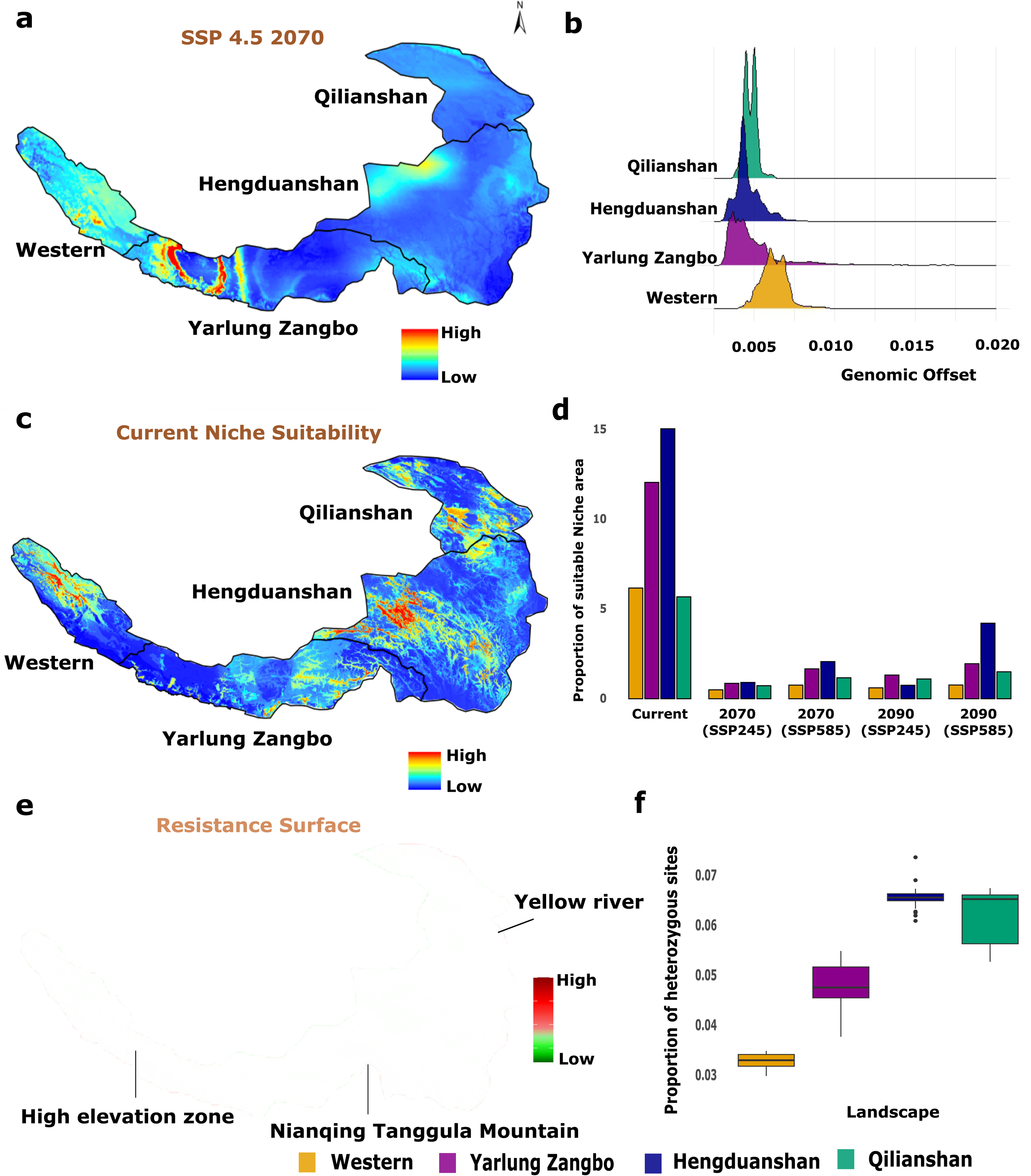
Climate change vulnerabilities in Tibetan Partridge populations revealed by landscape-scale genomic offset and niche modeling across the Sino-Himalayan region. **(a)** Genomic offsets predicted by the gradientForest model under SSP245 scenarios for 2061– 2080 across the Tibetan Partridge distribution range **(b)** Genomic offsets are significantly higher in the Western (n = 267,215 points), Yarlung Zangbo (n = 564,031 points), and Qilianshan (n = 1,030,066 points) landscapes, while the Hengduanshan landscape (n = 317,063 points) shows the lowest genomic offset values (p < 0.0001 for all pairwise comparisons, two-tailed Wilcoxon rank-sum test with FDR correction) **(c)** Current niche suitability model projected across the Tibetan Partridge’s distribution range **(d)** Proportions of suitable niche areas in each landscape under current and future climatic scenarios (SSP4.5 for 2070/2090 and SSP8.5 for 2070/2090) **(e)** Resistance surface map showing higher landscape connectivity (green) in the Hengduanshan and Qilianshan landscapes in the northeast compared to reduced connectivity in the Yarlung Zangbo and Western landscapes **(f)** Proportion of heterozygous sites in local Tibetan Partridge populations, indicating reduced heterozygosity in the Western landscape.

In addition to the population-scale genomic data, we also utilized the current distribution of the Tibetan partridge to quantify its present niche and predict future changes in the Sino-Himalayan landscape. We downloaded, and cleaned the distribution data of Tibetan Partridge from GBIF (https://www.gbif.org/) and performed ecological niche modeling using biomod2 (Thuiller et al. 2016). We combined the IUCN range map, and GBIF distribution data to delineate the distribution area of Tibetan partridge. Based on the distinct genetic clustering of the local Tibetan Partridge populations **(Fig. 2)** and the variable climatic environment they are exposed to **(Fig. 1)**, we modeled separately for two landscapes (western and north-eastern). Niche modeling analysis revealed significant variation in the bioclimatic variables that determined the suitability of habitats for Tibetan partridge populations across different landscapes **(Supplementary Table 3)**. In the western landscape, five bioclimatic variables; three related to temperature (BIO2, BIO4, BIO8) and two related to precipitation (BIO15, BIO19) had higher variable importance and therefore were important predictors of niche suitability **(Supplementary Table 3).** In contrast, in the north-eastern landscape, only three variables (BIO5; related to temperature, and BIO15/BIO14; related to precipitation) had higher variable importance, indicating diverse responses of different populations to their local climatic conditions.

Niche modeling analysis also revealed that 38.9% of the current habitat of Tibetan partridges is suitable, with the Hengduanshan landscape containing the largest proportion of suitable area (15.1%), followed by Yarlung Zangbo (12.05%) while the western (6.1%) and Qilianshan (5.6%) regions have the lowest suitability **(Fig. 4c, d).** This result indicate that there is differences in total suitable areas among landscapes in line with known pattern that species niche suitability declines towards the extreme of their distribution(Brown 1984). Under future projected climate change scenarios, the proportion of suitable areas significantly decreases across all landscapes **(Fig. 4d)**. Dry regions, particularly in the western landscape and outer areas of the Qilianshan landscape, were predicted to experience a significant decline in habitat suitability for Tibetan partridges. To assess the connectivity of the landscape for the potential movement of a species across its distribution range, we further used a simplified approach that combines the habitat suitability index with landscape variables such as elevation, elevation variability, and slope to create a resistance surface (Chen et al. 2022) (see method for details). Consistent with our EEMS findings **(Fig. 2f),** high resistance to species movement was observed across the Nianqing Tanggula mountains **(Fig. 4e)**, indicating limited dispersal ability between populations in western and north-eastern regions. Increased resistance was also identified along the edges of the Western and Yarlung Zangbo landscapes. In contrast, the Hengduanshan landscape displayed greater connectivity both within its area and with the Qilianshan landscape, suggesting stronger connectivity across northeastern landscapes.

These results indicated that Tibetan Partridge populations in the Western landscape, where suitable habitat constitutes only a small proportion (∼6%), exhibit the highest genomic offset. Expansive desert areas and the complex topography characterized by tall mountains in this region limits their ability to migrate toward the more suitable Hengduanshan landscape. Additionally, we observed low genetic diversity within these populations **(Fig. 4f)**, suggesting reduced genetic variation, which may limit their evolutionary flexibility and increase their vulnerability to changing environmental conditions. This pattern aligns with findings in isolated populations in other species that struggle to adapt to future environmental changes(Ruegg et al. 2018). Our findings are also consistent with previous studies on mammals, birds and plants in this region (Du et al. 2020; Chen et al. 2022; Chen et al. 2023; Jiao et al. 2024), which highlight how high-elevation mountains obstruct humid air, create drier conditions, restrict gene flow, and drive population divergence.

The Tibetan Partridge population in the Qilianshan landscape, located at the north-eastern edge of its range, shows a moderate genomic offset and limited proportion of suitable habitat (∼5%), identifying them as another potentially vulnerable populations. However, unlike the Western populations, this group may benefit from relatively better landscape connectivity and mid-elevation corridors, which could facilitate migration toward the more favorable Hengduanshan region. Additionally, the Qilanshan population also demonstrates higher genetic variation and admixture with the Hengduanshan population, suggesting greater genetic diversity could enhance its adaptability to environmental shifts. In contrast, the Yarlung Zangbo population, while sharing lower genetic variation and a moderate genomic offset with the Western population, benefits from a relatively higher proportion of suitable habitat (∼12%). Notably, this region harbors the lowest elevation Tibetan Partridge population at 2,900 meters, which diverged approximately 150,000 years ago during the Ghonge movement (Palacios et al. 2023). We believe this population has expanded its elevational niche, showcasing adaptations to diverse environmental conditions despite genetic constraints.

The Hengduanshan region, characterized by its complex terrain of intersecting mountain ranges and rivers, along with a diverse altitudinal range, fosters a diverse climatic environment. The local population of Tibetan Partridge from this region stands out for its high genetic diversity, a lower genomic offset, and a significant proportion of their habitat (∼15%) estimated as suitable for current and future niche availability. This region has been historically considered as a climate refugium(Sun et al. 2017), particularly during the Pleistocene, and has supported the survival of various bird and mammal species during past environmental shifts (Chen et al. 2022; Chen et al. 2023), and hence can be a crucial future climate refugium for Tibetan Partridges. Developing and rehabilitating movement corridors to facilitate gene flow between populations, could aid in enhancing the adaptability of Tibetan partridges for short and long-term environmental changes.

### Climate and biogeographic barriers shape evolutionary trajectories and population vulnerability in high-altitude species

The role of biogeographic barriers and climate in shaping species distributions, population divergence, and adaptation is a central question in ecology and evolutionary biology. While biogeographic influences are generally predictable, the interaction between climate and biogeography remains complex, particularly in the context of short and long-term environmental changes (De Kort et al. 2021). As global temperatures rise, precipitation patterns shift, and ecosystems transform, species must either adapt or face population decline and extinction. Examining divergence patterns and the mechanisms driving species’ responses helps us understand the interaction between climate and biogeography and whether species exhibit similar or contrasting adaptations across their range. Gaining this knowledge not only enhances our understanding of the mechanistic basis of species adaptation but also aids in identifying vulnerable populations, predicting their future changes, and developing effective conservation strategies.

This knowledge is particularly crucial for species inhabiting high-altitude environments, where complex geography isolates local populations and creates contrasting environmental conditions within relatively small landscapes (Sánchez-Montes et al. 2018). However, the logistical challenges of collecting samples in these regions often result in sparse sampling across environmental and geographic gradients, limiting our ability to draw clear and comprehensive conclusions. In this study, we successfully examined multiple local populations of a high-altitude endemic species across broad geographical and environmental gradients throughout its distribution range. This comprehensive sampling allowed us to examine the role of biogeographic features, such as mountains and valleys, and climate in shaping its population distribution patterns and mechanisms of local adaptation. By integrating high-throughput genomic data with landscape and ecological information, we have revealed contrasting climatic adaptations among populations across the species distribution range. Notably, we identify populations in drier regions as particularly vulnerable due to low genetic diversity, high genomic offsets, and reduced niche suitability under future climate change projections.

Our study provides valuable insights into the role of biogeography and climate shaping local adaptation of Tibetan Partridge populations across the Sino-Himalayan region, though these findings come with caveats. First, both predictive methods employed (genomic offset and niche modeling) have limitations (Costa et al. 2010; Rellstab et al. 2021). Genomic offset, as a relatively new concept, still requires more experimental validation (Rellstab et al. 2021).

Moreover, the data supporting our predictions rely on genomic and environmental metrics without comprehensive information on actual population sizes. Additionally, our niche modeling was constrained by species distribution data, especially in the western range, which may affect the accuracy of ecological projections. While climate data served as a basis for niche predictions, other factors such as habitat connectivity and biotic interactions like predation and competition are likely also influential in shaping the estimated niche of Tibetan Partridge (Wisz et al. 2013). Despite these constraints, this study contributes valuable insights into historical events, current local adaptations, and future vulnerabilities of Tibetan Partridges in the context of short and long-term environmental changes.

In summary, species inhabiting diverse environmental and climatic gradient like those in the Sino-Himalayan region, characterized by significant climatic fluctuations, face rapid environmental changes that challenge their survival. Populations with low genetic diversity, limited suitable habitat, high genomic offset, and poor habitat connectivity are particularly at risk **(Fig. 5).** To address these challenges, generating robust data on population history, genetic variation associated with local adaptation, and identification of vulnerable habitats and populations are critical for species conservation. This study highlights the value of integrating morphological, ecological, and evolutionary perspectives to uncover comprehensive insights into species’ historical dynamics, status, and future trajectories. Such an integrative approach is essential for designing and implementing effective conservation strategies that ensure the survival of species in the face of rapid environmental change.

**Figure 5:**
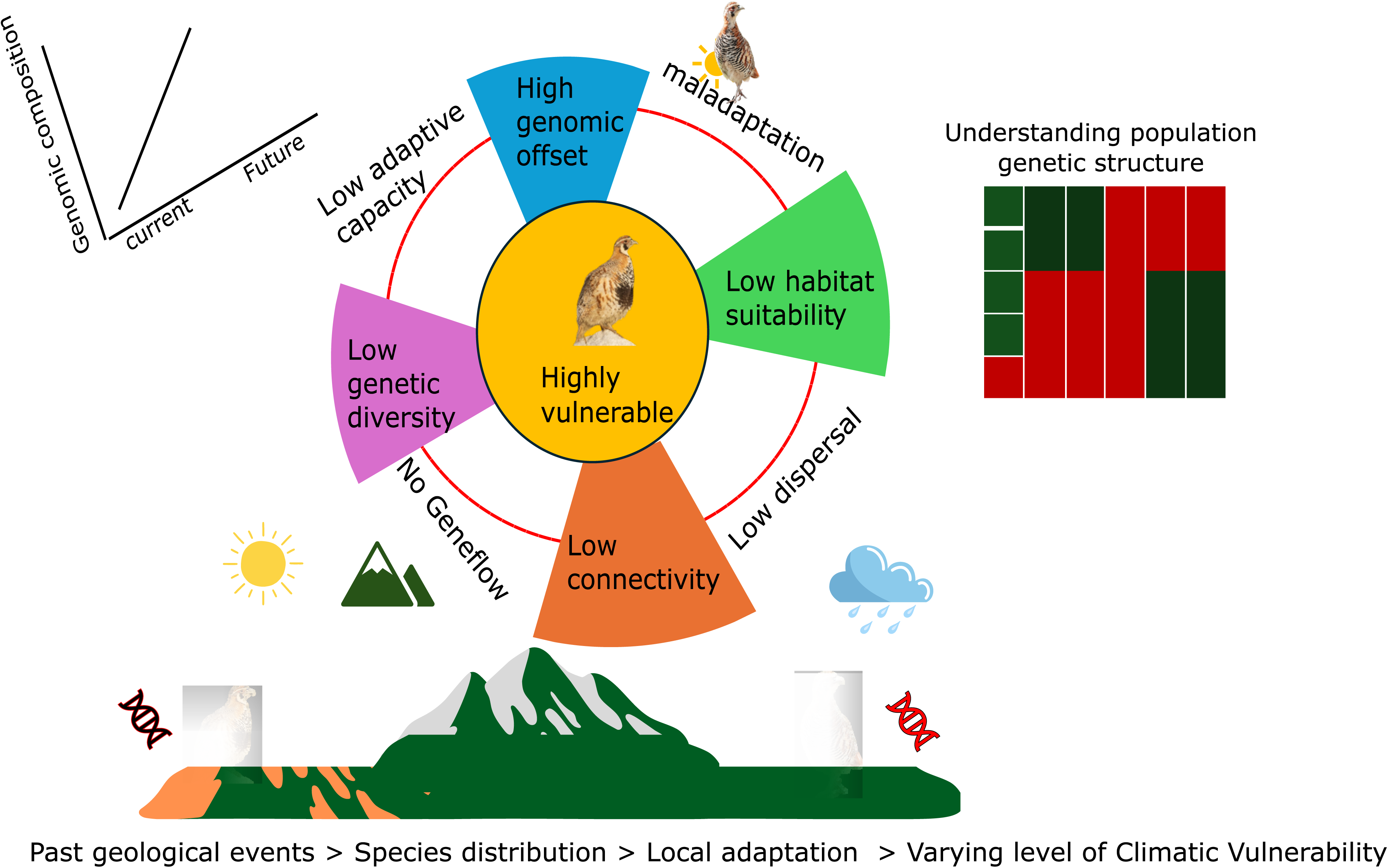
Conceptual framework of using genomic, ecological, and landscape information to understand climatic vulnerability. Past geological events shape the distribution of species such that the formation of barriers and subsequent local adaptation promote population divergence. These populations are exposed to varying levels of climatic vulnerability depending upon their genetics, ecology, and landscape connectivity.

## Materials and Methods

### Sample collection and genome re-sequencing

Blood samples were collected from 96 individuals of Tibetan Partridge across 28 different locations within the Qinghai-Tibetan Plateau (QTP) of the Sino-Himalayan landscape **(Fig. 1a)**. Eight of these populations are located to the west of the Yarlung Zangbo river and associated mountain ranges, while 20 are from the north-eastern regions, ensuring uniform sampling coverage across the distribution range of Tibetan Partridges.

Genomic DNA was extracted from samples using Qiagen’s DNeasy Blood and Tissue kit (Qiagen, Valencia, CA, USA). Whole-genome re-sequencing was performed for each individual using the Illumina NovaSeq 6000 platform, and the raw sequencing reads were mapped against the previously published genome of Tibetan Partridge (Palacios et al. 2023) using BWA v0.7.17 with default settings (Li and Durbin 2009). Quality control of the mapped reads was conducted using PICARD (http://picard.sourceforge.net/) to remove PCR duplicates, followed by the Genome Analysis Toolkit (GATK) v.4.2.0.0(McKenna et al. 2010) and GATK best practices recommendations (Van der Auwera et al. 2013) to perform base quality recalibrations, insertion/deletion (INDEL) realignment, Single Nucleotide Polymorphism (SNP) discovery, and genotyping across all 96 samples. The variant filtering was done using a pipeline previously developed in the lab (Lamichhaney et al. 2015), applying the following filtering parameters: Fisher strand bias > 15, mapping quality < 50, mapping quality rank sum < -0.1, read position rank sum < -3, base quality rank sum < -5, depth > 4000 and < 500, quality by depth < 5, SNP quality > 100, and genotype quality > 10. Variants genotyped across all 96 samples were retained, and the final VCF file was generated using VCFTools v.0.1.13 (Danecek et al. 2011) for downstream analyses.

### Morphological data collection

We measured morphological traits, including body length, tail length, and wing length using a flexible ruler. Beak length, head length, beak width, tarsus length, and middle toe claw length were measured with a vernier caliper, and body weight was recorded using an electronic scale. To account for body size variations, we adjusted the morphological data for body weight. These adjusted trait values were then used to assess differences along the landscape through boxplots, Analysis of Variance (ANOVA), and principal component analysis (PCA).

### Population genomics analysis

We used a filtered set of 5,068,604 high-quality SNPs to assess genetic diversity by computing population genetics parameters such as heterozygosity, and inter-population genetic diversity (FST) using VCFtools v.0.1.13 (Danecek et al. 2011). To identify phylogenetic relationships between samples we generated a maximum-likelihood phylogeny using 93,160 SNPs in FastTree v.0.2.1 with recommended default parameters for nucleotide alignments(Price et al. 2010). We further used SNAPP(Bryant et al. 2012) to estimate divergence time between seven phylogenetic groups (two outgroups: Daurian partridge and Grey Partridge, and individuals from six populations of Tibetan partridge across four different landscapes **(Fig. 1a)**, i. Western ii. Yarlung Zangbo high elevation iii. Yarlung Zangbo, 2900m elevation iv. Hengduanshan (Nanjiang valley) v. Hengduanshan (Sichuan) vi. Qilianshan, based on phylogenetic tree structure and known geographical barriers. We used Yarlung Zangbo’s 2900 m population and two populations from Hengduanshan (Sichuan and Nanjiang) to build a divergence timeline of different populations. We used the divergence time of genus *Perdix* genus diverged at 4.37 MYA (http://www.timetree.org/) to constraint with 0 offsets and a standard deviation of 0.005 million years. We ran BEAST v.2.6.3 (Bouckaert et al. 2019) for 5 million generations, with sampling conducted every 250 generations; the initial 10% was discarded as burn-in. We ran Tracer 1.7.2 (Rambaut et al. 2018) to ensure convergence of all parameters (ESS > 200) and TreeAnnotator 2.6.4 (Drummond and Rambaut 2007) to generate the maximum clade credibility tree.

We also estimated ancestry using a maximum-likelihood approach in Admixture v.1.3.0(Alexander et al. 2009) and plotted the admixture patterns across the landscapes using Mapmixture v.0.1.0(Jenkins 2024) in R. The optimal number of genetic ancestries was selected using a cross-validation procedure with the K-means method implemented in Admixture. To examine the pattern of isolation by distance (IBD) and isolation by environment (IBE), we performed the multiple regression for distance matrices (MRM) with 999 permutationst using genetic distance (FST), geographic distance (km), and Euclidean distance of selected environmental variables (BIO2, BIO8, and BIO9, BIO12, BIO15, see GEA below for details) using the ecodist package(Lichstein 2007) in R.

### Genotype Environment Association (GEA) analysis

We employed a combination of two independent approaches to identify genomic regions associated with bioclimatic variables. First, we utilized the Latent Factor Mixed Model (LFMM), a univariate analysis method that detects associations between allele frequencies and climatic variables (Frichot et al. 2013). To account for population structure in the dataset, we incorporated four latent factors identified by admixture analysis. We conducted five independent Markov Chain Monte Carlo (MCMC) runs, with a burn-in period of 5,000 iterations followed by 10,000 iterations. The results from these five runs were averaged and adjusted for multiple testing, applying a false discovery rate (FDR) correction of 1% as the significance threshold to identify candidate SNPs. To further validate and ensure the accurate identification of candidate SNPs, we calculated the Fixation index (FST) between two groups (five individuals each exposed to high and low values for each bioclimatic variable) using the candidate SNPs generated by LFMM alongside a comparable number of randomly selected SNPs across the genome.

The environmental variables for the multivariate analysis were carefully chosen based on correlation analysis and their importance ranking derived from gradient forest analysis, implemented using the R package "gradientForest"(Ellis et al. 2012). This approach ensured the inclusion of relevant variables while minimizing redundancy in the analysis. Due to computational limitations, we only used ∼500,000 SNPs from the longest scaffold (Scaffold 738) for this analysis. The selected predictors included three temperature-related variables: BIO2 (Mean Diurnal Range, calculated as the mean of monthly maximum temperature minus minimum temperature), BIO8 (Mean Temperature of Wettest Quarter), and BIO9 (Mean Temperature of Driest Quarter), as well as two precipitation-related variables: BIO12 (Annual Precipitation) and BIO15 (Precipitation Seasonality, measured as the Coefficient of Variation) **(Supplementary Fig. 3)**.

Next, we employed Redundancy Analysis (RDA), a multivariate approach, to examine the relationship between multiple environmental variables and SNPs using the vegan package in R(Capblancq and Forester 2021; Oksanen et al. 2024). Genetic variations associated with the environment were considered significant if they had loadings in the tails of the distribution, specifically within a standard deviation limit of 3.5 along the first three RDA axes, as recommended in(Forester et al. 2018).

To perform functional analysis of the candidate genomic regions, we annotated the filtered VCF file using the published genome of Tibetan Partridge(Palacios et al. 2023) using SnpEFF (Cingolani et al. 2012). We then conducted Gene Ontology (GO) term analysis to identify enriched GO terms among the candidate SNPs, comparing them against the background annotated gene universe in the reference genome in R. First, we annotated the Tibetan partridge transcriptome previously published(Palacios et al. 2023) using eggNOG-mapper (Cantalapiedra et al. 2021) with the default parameters for characterizing Gene Ontology (GO) terms. We then constructed an organism database (OrgDB) library for Tibetan partridge using the makeOrgPackage function of AnnotationForge package (Carlson and Pagès 2024) in R. Gene Ids of the climate-associated candidate SNPs were extracted and GO enrichment analysis was done using the enrichGO and enricher functions from tclusterProfiler (Wu et al. 2021) package in R. We considered GO terms with adjusted p-value and q-value < 0.05 as statistically significant enrichment.

### Genomic offset prediction

We used a machine learning-based non-parametric modeling approach to predict allelic turnover under future climatic conditions, using the R package gradientForest(Ellis et al. 2012). This method calculates genetic offset as the Euclidean distance between genomic compositions in the current and predicted future climatic scenarios. To carry out this analysis, we downloaded three general circulation models (GCMs)— FIO-ESM-2, IPSL-CM6A-LR, and UKESM1-0-LL, from the Worldclim database (http://www.worldclim.org) (Fick and Hijmans 2017). These models correspond to two carbon emission scenarios based on shared socioeconomic pathways (SSPs): SSP 4.5 (moderate) and SSP 8.5 (harsh), for two time periods: 2061-2080 and 2081-2100, all at a resolution of 30 seconds. For each given GCM, time period, and SSPs, we built GF models (12 models) and estimated the genetic offset for the same five bioclimatic variables previously used in Gene-Environment Analysis. We averaged the predictions for each SSPs across the two time periods, yielding four overall predictions (SSPs 4.5-year 2070, SSPs 8.5 year 2070, SSPs 4.5 year 2090, and SSPs 8.5 year 2090). We used the Wilcoxon rank test to quantify the pairwise statistical difference of genomic offset between four landscapes.

### Niche modeling

To assess the current climatic niche and predict future suitability for the Tibetan Partridge, we utilized occurrence data of Tibetan Partridge collected from field sampling and the GBIF database (GBIF 2023). After filtering occurrences at a 1-km resolution, we retained a total of 230 data points, with 96 occurrences from the eastern range and 134 from the western range. We modeled these two ranges separately to identify bioclimatic variables that serve as significant predictors of niche suitability. We used the same bioclimatic data downloaded for genomic offset predictions used above. We further employed the USDM (version 2.1) package (Naimi et al. 2014) in R to assess the variance inflation factor (VIF) and identified multicollinearity among predictor variables, removing any variables with a VIF greater than 2.5. Additionally, we evaluated the variable importance using the Biomod2 package (Thuiller et al. 2016), retaining only those with higher variable importance.

We further utilized an ensemble modeling approach within the Biomod2 package (Thuiller et al. 2016) in R to assess species distribution under both current and future climatic scenarios. For modeling, we randomly sampled 1,000 background points for both the western and north-eastern landscapes. We applied three iterations of the cross-validation procedure, randomly dividing occurrence records into two subsets: 70% of the data for model calibration and the remaining 30% for testing. Model evaluation was based on the Receiver Operating Characteristic (ROC), a threshold-independent indicator of model performance (Hanley and McNeil 1982), and the True Skill Statistic (TSS), which is calculated as the sum of sensitivity and specificity minus one (Allouche et al. 2006). The predictions from various modeling algorithms were summarized using a weighted mean approach (Marmion et al. 2009), excluding models with a TSS of 0.60 or lower in the final model construction.

### Landscape resistance surface analysis

We employed a simplistic approach to map the resistance surface across the Tibetan Partridge’s distribution range, drawing from the methodology of a previous study (Chen et al. 2022). To calculate the resistance surface, we first computed the average values of key landscape features (elevation, standard deviation of elevation, and slope) using the current distribution of Tibetan Partridges as a reference. For each grid in the study area, we calculated the absolute difference between the value of each landscape feature and the average value of that feature based on the current distribution. A smaller difference indicated lower resistance, while a larger difference suggested greater resistance to movement. Next, we incorporated habitat suitability information into the resistance surface by subtracting the value of each grid from the highest suitability value, where a lower resultant value indicated lower resistance. To make the resistance surface easier to visualize and interpret, we rescaled the average of these continuous landscape values and habitat suitability to a scale from 1 (lowest resistance) to 10 (highest resistance) (Chen et al. 2022).

## Acknowledgements

The fieldwork and genome sequencing were supported by the Central Forestry and Grassland Ecological Protection and Restoration Fund in Qinghai Province, the National Survey on Terrestrial Wildlife Resources in China, and The Abundance, Distribution, and Habitat of Leopards and Their Main Preys in Eastern Xizang (LC-3-04) to NW. PG was supported by a graduate assistantship from Kent State University. We express our gratitude to the personnel at the National Forestry and Grassland Administration of China, and the Forestry and Grassland Administration of Qinghai, Tibetan, and Sichuan provinces, China, for their unwavering support throughout the project. Special thanks to Hongtao Mao, Honglei Li, Qinze Zhang, Jiahu Jiang, Yichuan Meng, and Rong Liu for their contribution to the fieldwork associated with this study. We are thankful to Dr. Kristen Ruegg and Dr. Joan Ferrer Obiol for their insightful comments on the first draft of the manuscript. We also thank Neeraja V, Nick Addey, and Macauley Library at Cornell Lab of Ornithology for permitting us to use their photographs in this paper.

## Supplementary Figures legends

**Supplementary Figure 1: Morphological trait variation among Tibetan Partridge populations residing across different landscapes (a)** Body length, **(b)** Left paw length, **(c)** Right paw length, **(d)** Tail length, **(e)** Left-wing length, **(f)** Right-wing length, **(g)** Left tarsometatarsus length, and **(h)** Right tarsometatarsus length. P-values displayed are derived from Analysis of Variance (ANOVA), with p < 0.05 denoting significant differences in trait values across landscapes at the 95% confidence interval. Adjusted value for each trait calculated by dividing raw measure by body weight.

**Supplementary Figure 2: Topographic and vegetational heterogeneity in the Sino-Himalayan region (a)** 3-dimensional elevation map highlighting significant elevation variability across the landscape, with high-elevation regions (in yellow) aligned with key topographic features indicated in Fig. 1, contrasting with the plateau areas in the northeast (green), which range from 4000m to 5000m. (map source: WorldviewR: https://jcallura.github.io/) **(b)** The Normalized Difference Vegetation Index (NDVI) map shows vegetation density differences across the distribution range. The northeast region exhibits higher vegetation cover, while western areas have relatively sparse vegetation. (Data source: MODIS 16-day vegetation index, 1 km resolution, for June 1, 2020, processed using MODIStsp in R(Busetto and Ranghetti 2016).

**Supplementary Figure 3: Variation in sequencing coverage of individual samples across landscape.** Gray dot refers to each individual sample’s coverage.

**Supplementary Figure 4: Molecular dating and genetic divergence in Tibetan Partridge populations (a)** Molecular dating analysis indicates recent divergence within Tibetan partridge populations, with divergence times ranging from 0.33 to 0.05 million years ago (mya). The north-eastern and western populations, separated by the Nianqing Tanggula mountains, diverged around 0.33 mya. The population along the Nujiang River in the Hengduanshan landscape, northeast of the Nianqing Tanggula mountains, diverged from the plateau population approximately 0.10 mya. The Yarlung Zangbo and Western populations, separated by high-elevation zones along the upper Yarlung Zangbo river, diverged around 0.05 mya; the most recent divergence within Tibetan partridges is low elevation movement to 2900 m. Genetic variation across landscapes can be poorly explained by (b) Isolation by Distance (IBD), though its effects are relatively weak((β = 0.0000433, p = 0.125). In contrast, (c) Isolation by Environment (IBE) has a stronger and more significant influence β = 0.0771, p = 0.001)

**Supplementary Figure 5: Gradient forest analysis highlights precipitation-related variables (blue) as the strongest predictors of genetic variation in Tibetan Partridges (a)** Ranked importance of variables based on prediction accuracy and **(b)** R-squared weighted importance reveals the dominance of precipitation-related factors. Variables in grey were excluded from further modeling to prioritize climate-related predictors.

**Supplementary Figure 6: Manhattan plot showing -log10(p) values from Gene Environment Association (GEA) analysis to identify association with bioclimatic variables (a)** BIO2 (Mean Diurnal Range) **(b)** BIO8 (Mean Temperature of Wettest Quarter) and **(c)** BIO15 (Precipitation Seasonality). SNPs above the red dashed line denote candidate SNPs after a significance threshold of 1% False Discovery Rate (FDR) correction.

**Supplementary Figure 7: Distribution of genetic differentiation (FST) for “climate-associated core” candidate SNPs vs. random SNPs,** for **(a)** BIO2 (Mean Diurnal Range) **(b)** BIO8 (Mean Temperature of Wettest Quarter) **(c)** BIO9 (Mean Temperature of Driest Quarter). **(d)** BIO12 (Annual Precipitation) and **(e)** BIO15 (Precipitation Seasonality). FST calculations were based on ten individuals each from populations exposed to the highest and lowest values for each respective bioclimatic variable.

**Supplementary Figure 8: Venn diagram illustrating the overlap of candidate SNPs identified by univariate (LFMM) and multivariate (RDA) analyses associated with bioclimatic variables;** center: overlapping SNPs identified by both LFMM and RDA, suggesting robust associations with climatic adaptation, lower left: number of SNPs identified by LFMM analysis for individual bioclimatic variables, with precipitation-related variables shown in green and temperature-related variables in red in the right table.

**Supplementary Figure 9: The gradientForest predicted genomic offsets (a)** under *SSP 8.5* for the year 2070 (p < 2e^-16^ for all pairwise comparisons, the two-tailed Wilcoxon rank-sum test and FDR correction for multiple comparisons) **(b)** under *SSP 4.5* for the year 2090 (p < 2e^-16^ for all pairwise comparison, the two-tailed Wilcoxon rank-sum test and FDR correction for multiple comparisons) and **(c)** under *SSP 8.5* for the year 2090 (p < 2e^-16^ for all pairwise comparison, the two-tailed Wilcoxon rank-sum test and FDR correction for multiple comparisons). *SSP 8.5* is the worst climatic conditions and *SSP 4.5* is moderate climatic conditions. The distribution of predicted genomic offset across each landscape is shown in the right panel for each figure.

## Data availability

All short-read whole-genome sequencing data have been submitted to the National Center for Biotechnology Information (NCBI) under BioProject Accession number PRJNA1198335.

## Code availability

Scripts and workflow for all analyses done in this paper is available on the GitHub page https://github.com/Prashantevo/Tibetanpatridgeeeb.

## Competing interests

The authors declare no competing interests.

## Additional information

Supplementary Materials

Supplementary Resources 1 - 2

## Notes

### Competing Interest Statement

The authors have declared no competing interest.

### Summary of Updates

This version of the manuscript has been revised to update the author name, which was mistaken in the previous version.

