## Supplementary Fig for "Integrated landscape genomics reveals biogeography and climate-driven local adaptation in high-altitudes"

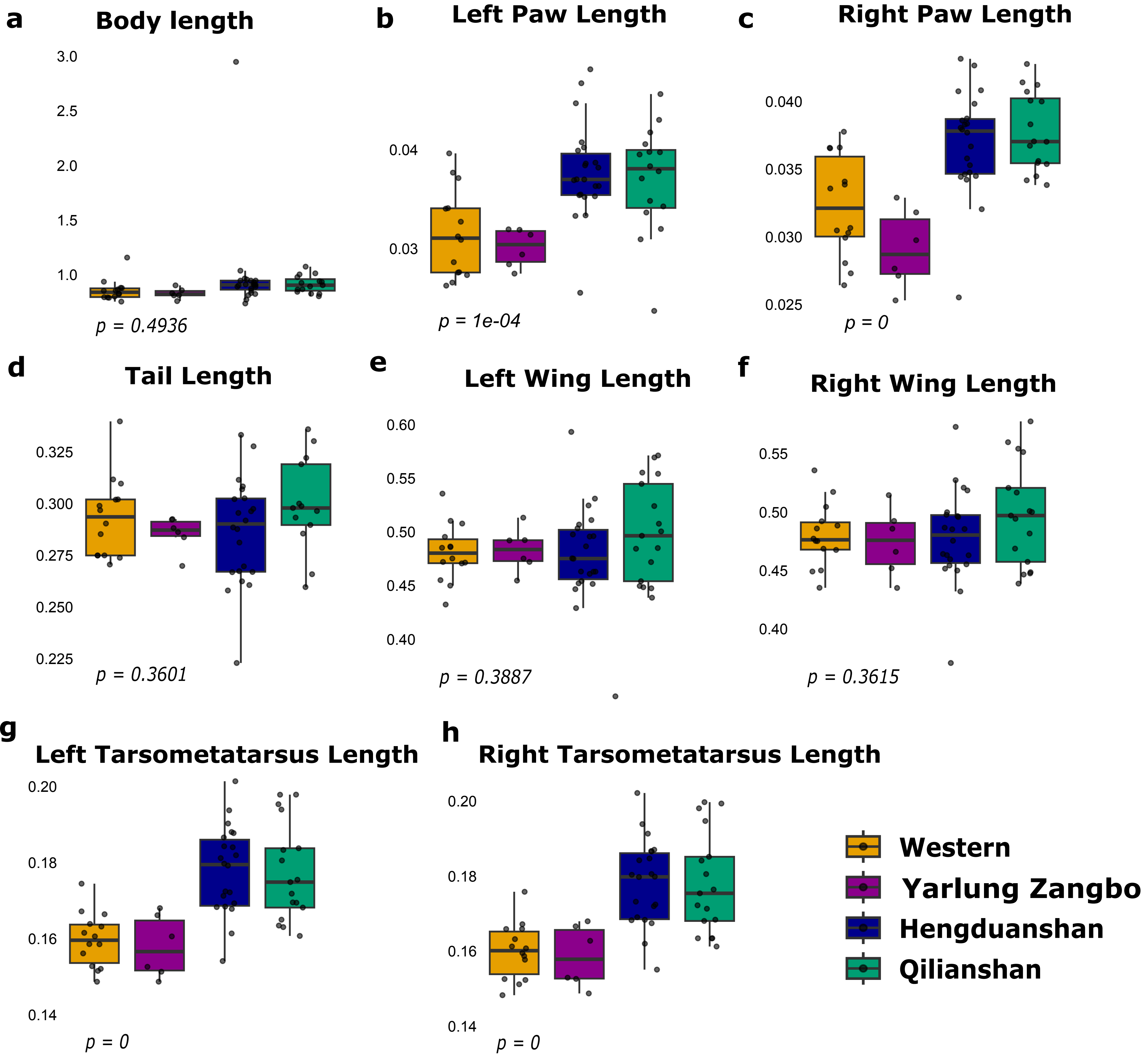


**Supplementary Figure 1: Morphological trait variation among Tibetan Partridge populations residing across different landscapes (a)** Body length, **(b)** Left paw length, **(c)** Right paw length, **(d)** Tail length, **(e)** Left wing length, **(f)** Right wing length, **(g)** Left tarsometatarsus length, and **(h)** Right tarsometatarsus length. p-values displayed are derived from Analysis of Variance (ANOVA), with $p < 0.05$ denoting significant differences in trait values across landscapes at the 95% confidence interval. Adjusted value for each trait calculated by dividing raw measure by body weight.

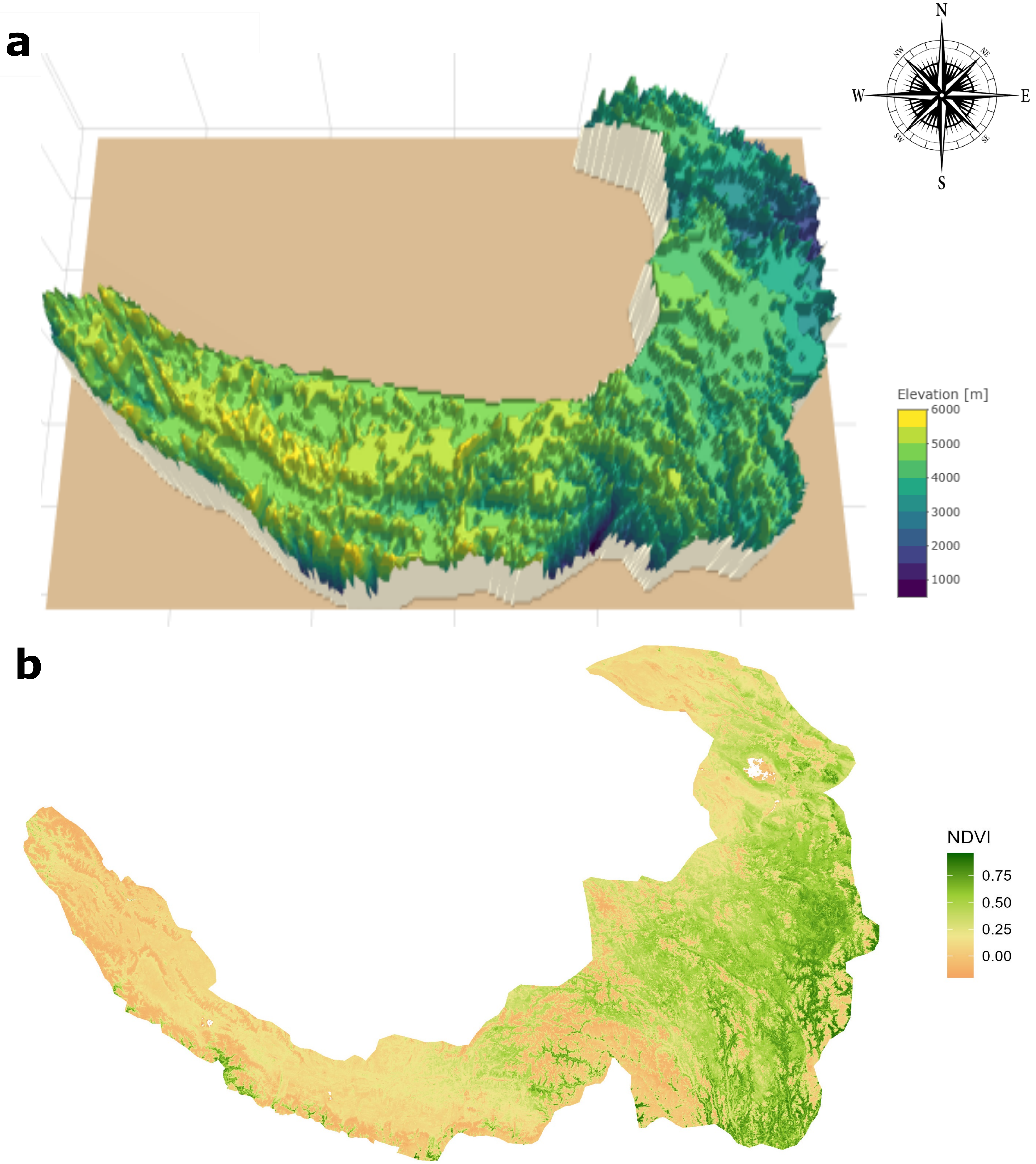


**Supplementary Figure 2: Topographic and vegetational heterogeneity in the Sino-Himalayan region (a**) 3-Dimensional elevation map highlighting significant elevation variability across the landscape, with high-elevation regions (in yellow) aligned with key topographic features indicated in Fig. 1, contrasting with the plateau areas in the northeast (green), which range from 4000m to 5000m. (map source: WorldviewR: https://jcallura.github.io/)
**(b)** The Normalized Difference Vegetation Index (NDVI) map shows vegetation density differences across the distribution range. The northeast region exhibits higher vegetation cover, while western areas have relatively sparse vegetation. (Data source: MODIS 16-day vegetation index, 1 km resolution, for June 1, 2020, processed using MODIStsp in R[73].

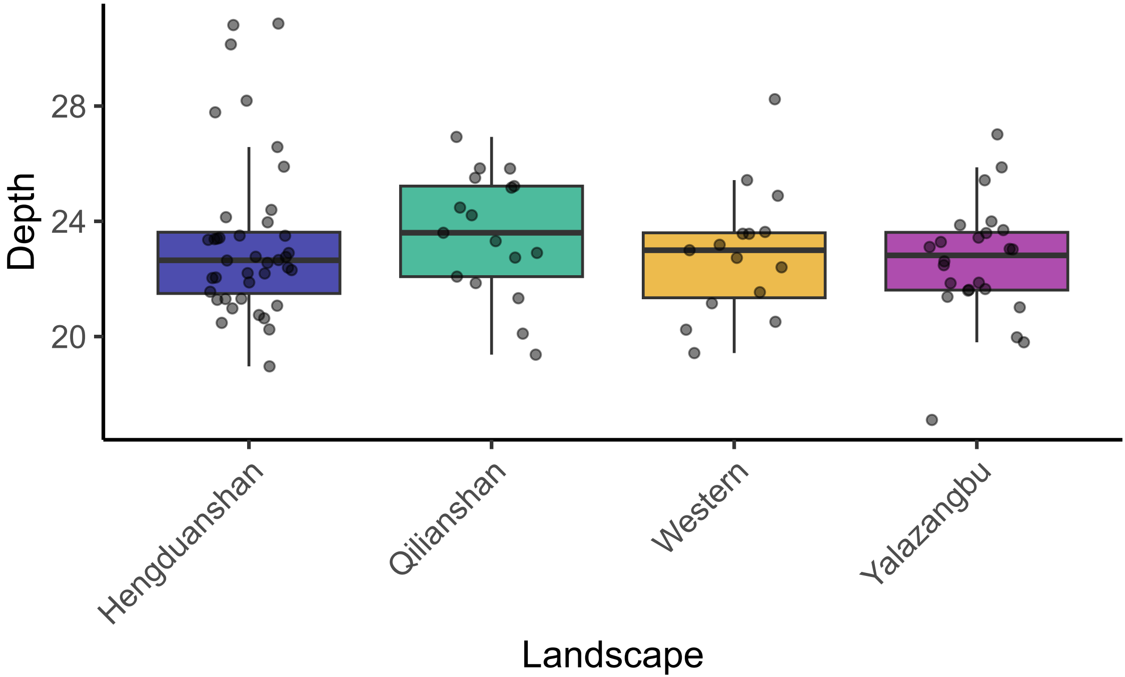


**Supplementary Figure 3: Variation in sequencing coverage of individual samples across landscape.** Gray dot referes to each individuals sample's coverage**.**

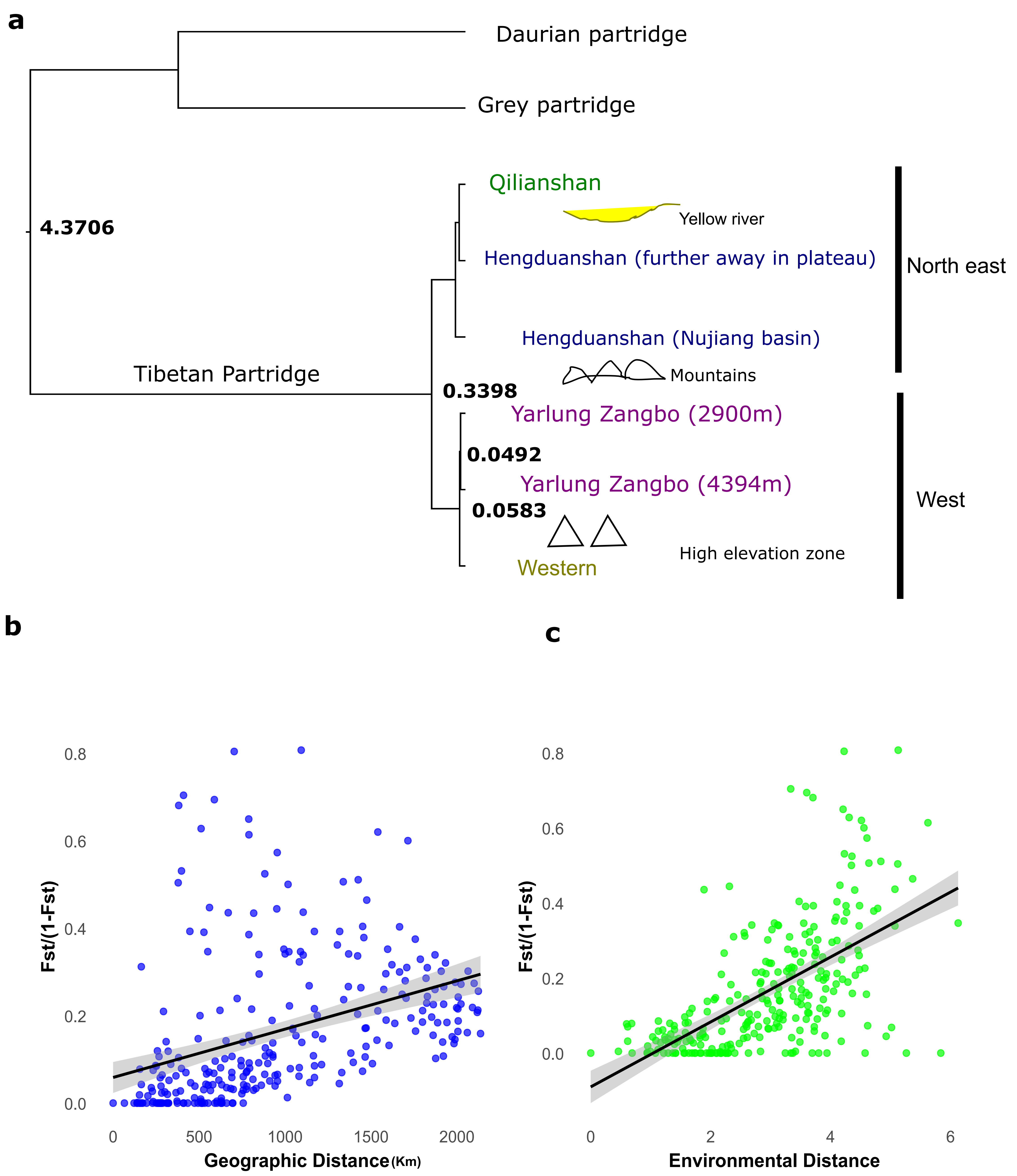


**Supplementary Figure 4: Molecular dating and genetic divergence in Tibetan Partridge populations**
**(a)** Molecular dating analysis indicates recent divergence within Tibetan partridge populations, with divergence times ranging from 0.33 to 0.05 million years ago (mya). The north-eastern and western populations, separated by the Nianqing Tanggula mountains, diverged around 0.33 mya. The population along the Nujiang river in Hengduanshan landscape, northeast of the Nianqing Tanggula mountains, diverged from the plateau population approximately 0.10 mya. The Yarlung Zangbo and Western populations, separated by high-elevation zones along the upper Yarlung Zangbo river, diverged around 0.05 mya; the most recent divergence within Tibetan partridges is low elevation movement to 2900 m. Genetic variation across landscapes can be poorly explained by (b) Isolation by Distance (IBD), though its effects are relatively weak ($\beta = 0.0000433$, $p = 0.125$). In contrast, (c) Isolation by Environment (IBE) has a stronger and more significant influence ($\beta = 0.0771$, $p = 0.001$).

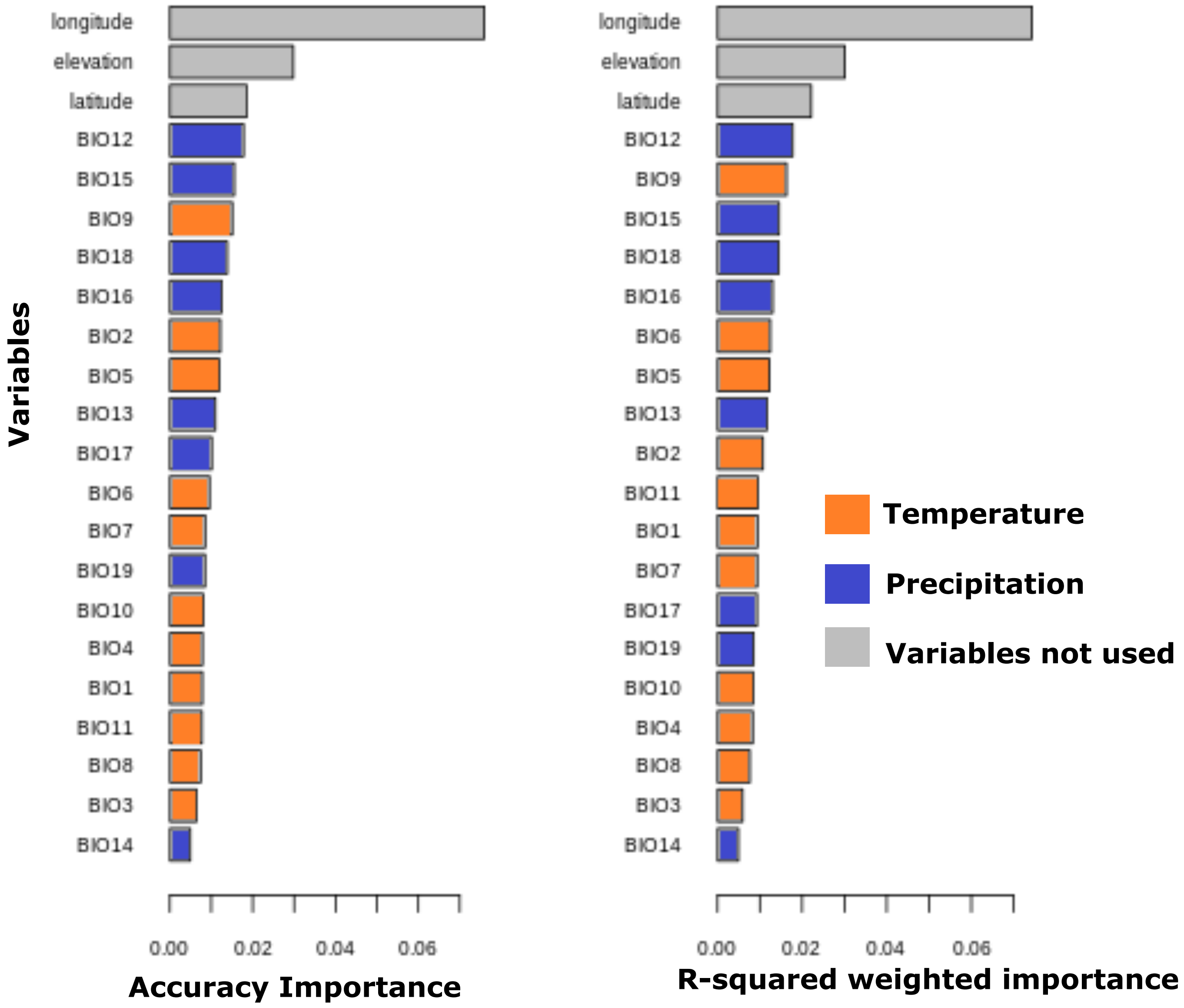


**Supplementary Figure 5: Gradient forest analysis highlights precipitation-related variables (blue) as the strongest predictors of genetic variation in Tibetan Partridges (a)** Ranked importance of variables based on prediction accuracy and **(b)** R-squared weighted importance reveal the dominance of precipitation-related factors. Variables in grey were excluded from further modeling to prioritize climate-related predictors.

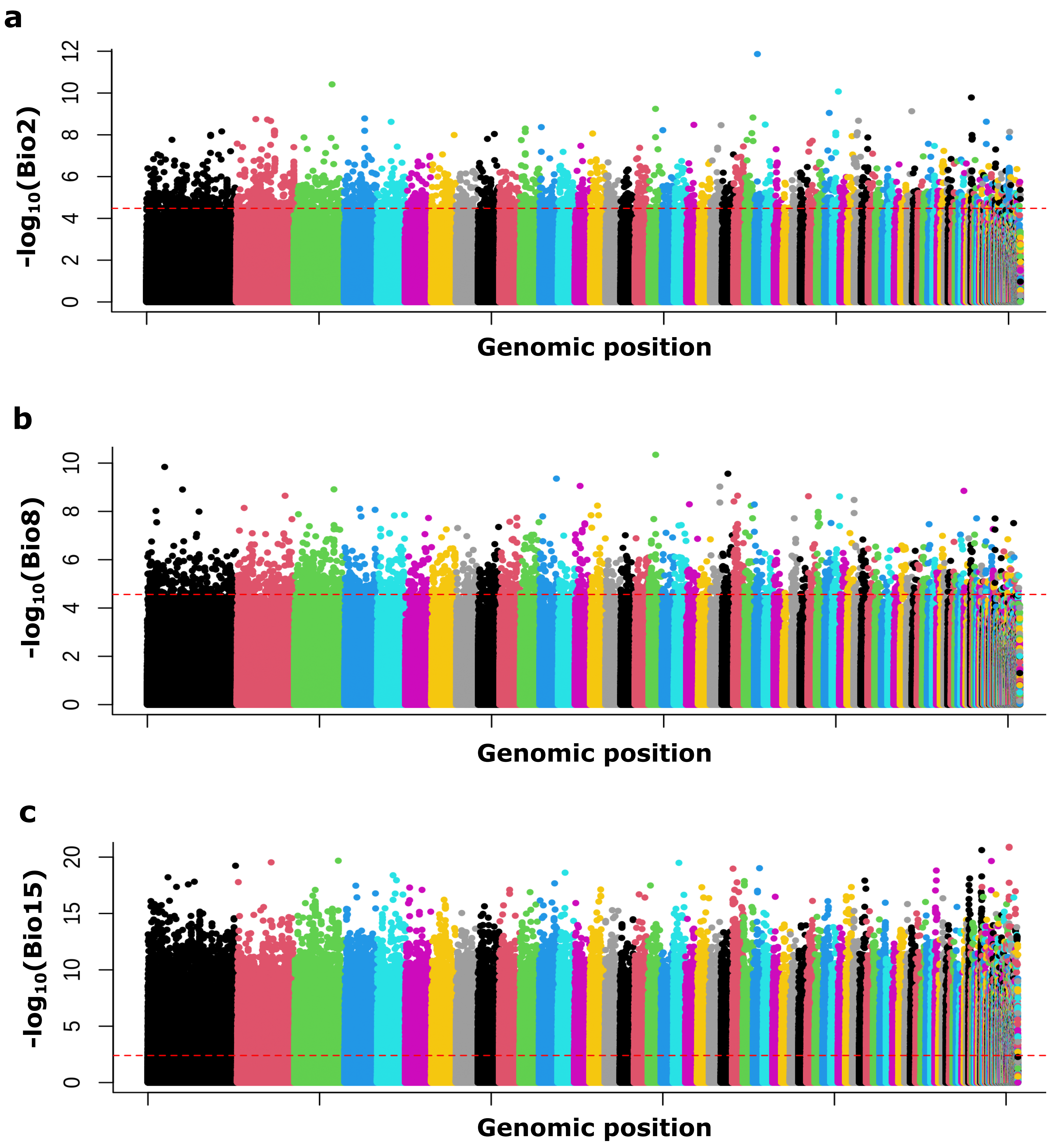


**Supplementary Figure 6: Manhattan plot showing -$\log_{10}$(p) values from Gene Environment Association (GEA) analysis to identify association with bioclimatic variables (a)** BIO2 (Mean Diurnal Range) **(b)** BIO8 (Mean Temperature of Wettest Quarter) and **(c)** BIO15 (Precipitation Seasonality). SNPs above red dashed line denotes candidate SNPs after significance threshold of 1% False Discovery Rate (FDR) correction.

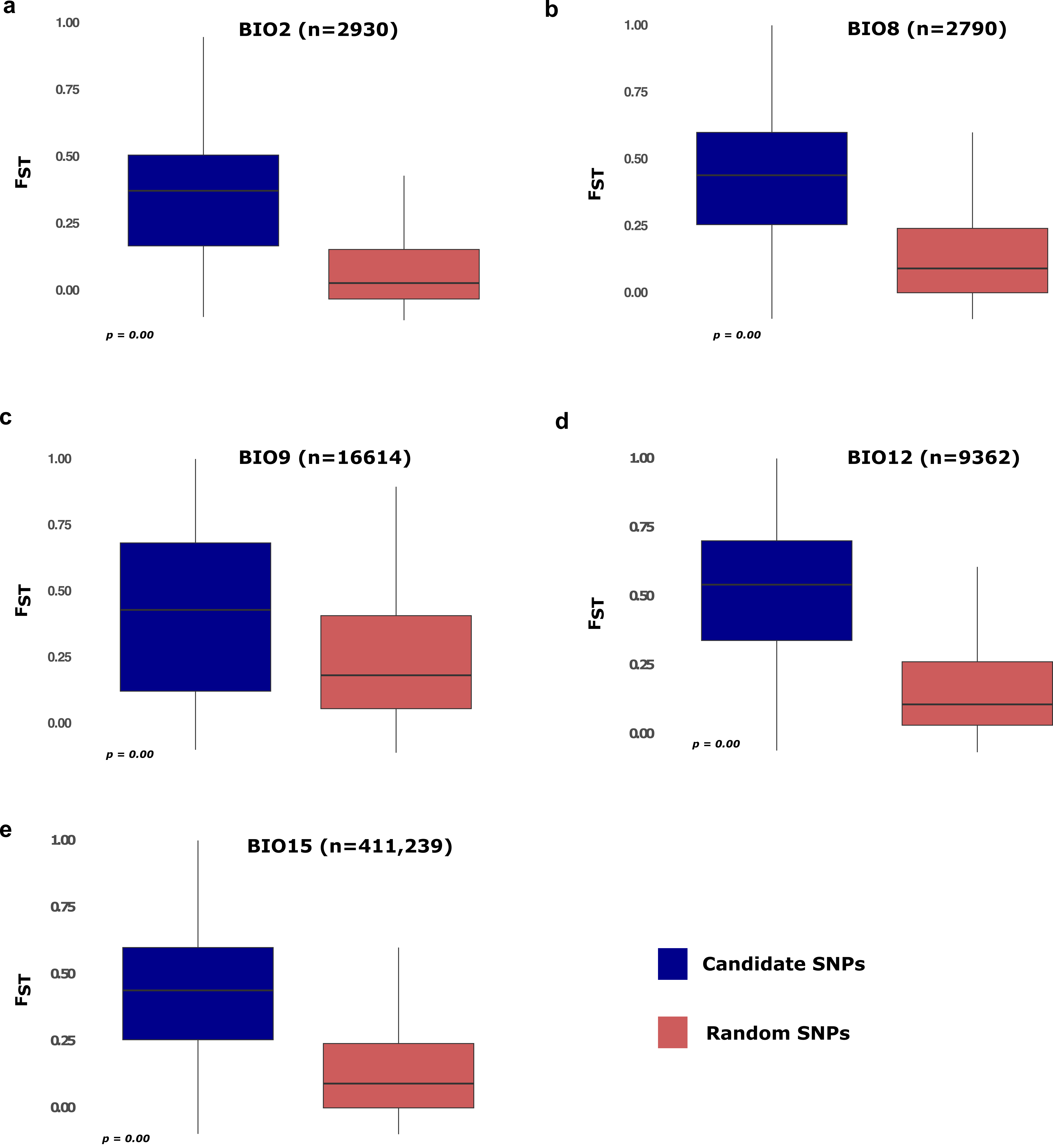


**Supplementary Figure 7: Distribution of genetic differentiation (FST) for "climate-associated core" candidate SNPs vs. random SNPs, for (a)** BIO2 (Mean Diurnal Range) **(b)** BIO8 (Mean Temperature of Wettest Quarter) **(c)** BIO9 (Mean Temperature of Driest Quarter) **(d)** BIO12 (Annual Precipitation) and **(e)** BIO15 (Precipitation Seasonality). FST calculations were based on ten individuals each from populations exposed to highest and lowest values for each respective bioclimatic variable.

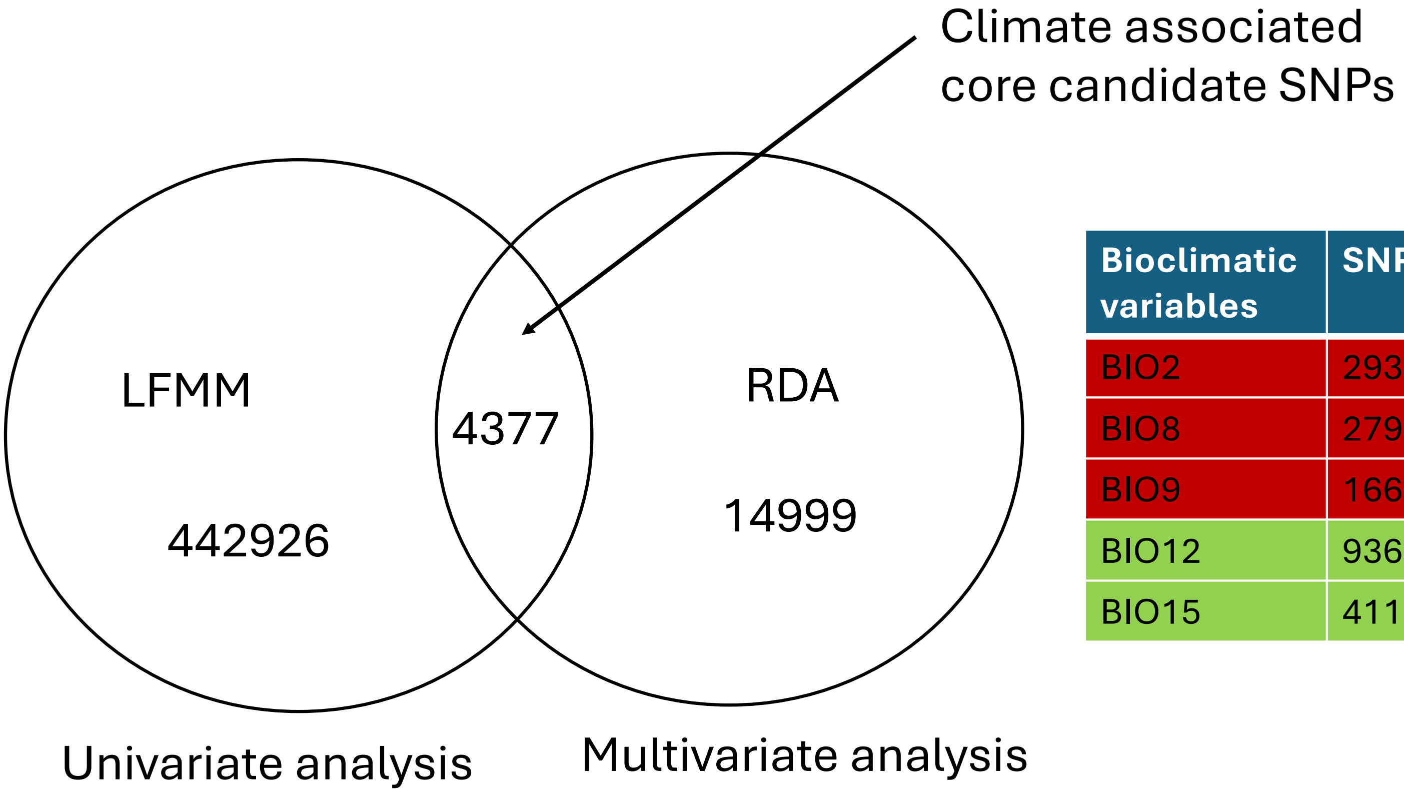


| Bioclimatic variables | SNPs |
|---|---|
| BIO2 | 2930 |
| BIO8 | 2790 |
| BIO9 | 16614 |
| BIO12 | 9362 |
| BIO15 | 411230 |

**Supplementary Figure 8: Venn diagram illustrating the overlap of candidate SNPs identified by univariate (LFMM) And multivariate (RDA) analyses associated with bioclimatic variables;** center: overlapping SNPs identified by both LFMM and RDA, suggesting robust associations with climatic adaptation, lower left: number of SNPs identified by LFMM analysis for individual bioclimatic variables, with precipitation-related variables shown in green and temperature-related variables in red in the right table.

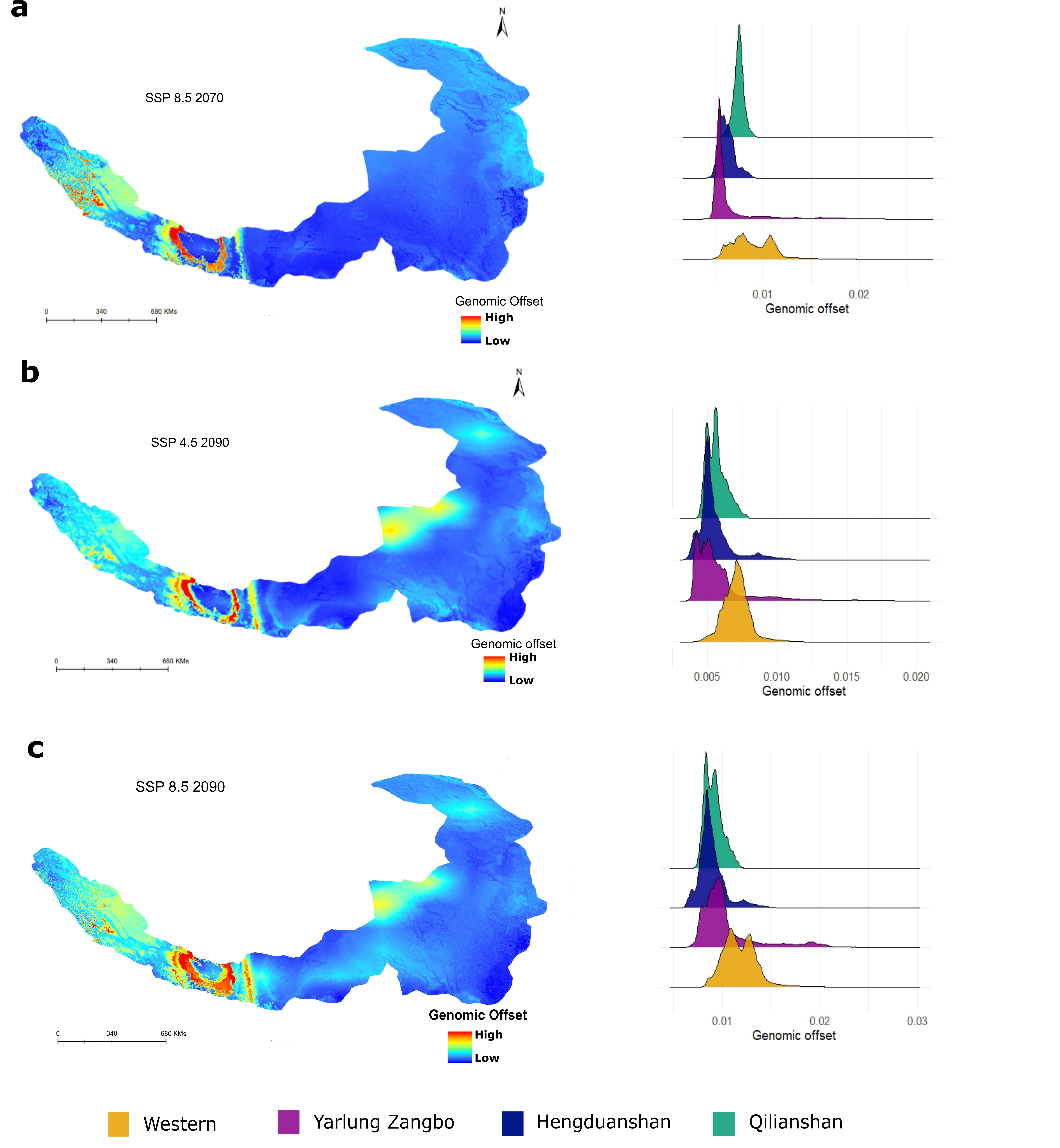


**Supplementary Figure 9: The gradientForest predicted genomic offsets (a)** under SSP 8.5 for the year 2070 ($p < 2e-16$ for all pairwise comparison, the two-tailed Wilcoxon rank-sum test and FDR correction for multiple comparisons) **(b)** under SSP 4.5 for the year 2090 ($p < 2e-16$ for all pairwise comparison, the two-tailed Wilcoxon rank-sum test and FDR correction for multiple comparisons) and **(c)** under SSP 8.5 for the year 2090 ($p < 2e-16$ for all pairwise comparison, the two-tailed Wilcoxon rank-sum test and FDR correction for multiple comparisons). SSP 8.5 is worst climatic conditions and SSP 4.5 is moderate climatic conditions. Distribution of predicted genomic offset across each landscape is shown in the right panel for each figure.
